# Optimal Cellular Phenotypic Adaptation in Fluctuating Environments

**DOI:** 10.1101/2023.01.17.524479

**Authors:** Jason T. George

## Abstract

Phenotypic adaptation is a core design feature of bacterial populations and multicellular systems navigating highly variable environments. Recent empirical data implicates the role of memory-driven decision-making in cellular systems navigating uncertain future nutrient environments based on prior experience, wherein a distinct growth phenotype emerges in fluctuating conditions. We develop a simple stochastic mathematical model to describe the cellular decision-making required for systems to optimally navigate such uncertainty. We demonstrate that adaptive populations capable of sensing their environment and estimating the nutrient landscape more efficiently traverse changing environments. We find during environmental transitions that larger memory capacities strike a trade-off between inertia of past environmental memory and higher resolution for estimating the optimal phenotype whenever the underlying landscape is close to a critical break-even point. Moreover, systems that tune their memory capacity avoid growth penalties resulting from maladaptive phenotypes following changes to the metabolic landscape. Our model predicts that the nutrient availability of adaptive cells is universally reduced in fluctuating nutritional environments relative to those in constant ones, which recapitulates empirical observations in bacterial systems. Our findings demonstrate that this deviation is a consequence of environmental mis-estimation together with bet-hedging in uncertain adaptive landscapes, and suggests that this deviation is fully determined by cellular memory capacity and the proximity of the environmental landscape to the system’s critical break-even environment. We anticipate that our mathematical framework will be more broadly useful for studying memory-driven cellular decision-making in biological contexts where there is a trade-off for cells selecting from multiple phenotypic states. Such a tool can be used for predicting the response of complex systems to environmental alterations and for testing therapeutically relevant policies.

## 1. Introduction

Biological systems commonly encounter and respond to exogenous environmental fluctuations, and their consequent adaptation generates a diversity of phenotypic responses at the population level [1–3]. In bacterial populations, dramatic phentypic changes resulting in bacterial persistence and longeterm infection can occur with relatively few molecular events in simple biological circuits[4–6]. Similar phenomena occur in mammalian systems and complicate therapeutic intervention with drug and targeted resistance [7–9].

Prior work has studied phenotypic switching in response to fluctuating environments in several biological contexts. Deterministic modeling characterized the tradeoff between stress resistance and growth-lag [10], where the mean behavior in response to stochastic fluctuations was considered by invoking the law of large numbers over sufficiently many cycles. In another study, combined modeling and experimental analysis explored the rate of variable phenotypic switching relative to that of environmental fluctuation [11]. These results predicted that cells matching their rate of phenotypic switching to the characteristic rate of environmental fluctuation exhibited enhanced growth. Recent models of phenotypic switching have also been applied to better characterize stochastic state transitions between stable local minima in gene regulatory networks [12].

Cellular memory is a key defining feature of adaptive cells in changing environments [13], and recent empirical evidence reveals a number of mechanisms by which this can occur at the single-cell level. Previous studies have shown that yeast cells with prior stress exposure and consequent adaptation-driven tolerance retain an associated memory across generations via transcriptional control of long-lived stress-related proteins [14]. Since then, additional molecular mechanisms of cellular memory have been identified: Durable chromatin modifications can persist during cellular division [15]. Prior growth-factor signaling induces short-term memory that tunes subsequent receptor sensitivity [16]. Even oligomeric protein condensates arising in response to previous environmental encounters enable cells to form memories that affect future function [17]. Recent work has begun to highlight the importance of memory-driven decision-making in navigating future events, not just present ones [18–20].

While these features are of direct importance for understanding cellular adaptation, the function of memory on stochastic optimal decision-making and its impact on cell fitness are at present unknown. To address this, and to explain recent empirical evidence of distinct reduced-growth phenotypes in fluctuating environments [21], we developed a simplified mathematical framework to model memory-driven optimal phenotypic switching in stochastic environments. In our model, adaptive populations leverage a memory of historical environmental exposures to forecast the optimal decision strategy for unknown future landscapes. This foundational model is amenable to explicit analytic characterization of the optimal strategies as functions of environmental history and memory capacity. In the setting of metabolic navigation, our model quantifies the dynamical differences between static cellular phenotypes committed to navigating a single environment and those that may select a phenotypic strategy based on their memory of past experience in order to optimize their growth potential. In studying the role of memory capacity, we identify a fundamental trade-off between the accuracy of a cell’s estimation of the current environmental state and an associated momentum delaying the rate of adaptation to new environments. We quantify the efficiency of systems employing a dynamic memory scheme that balance the above constraints when compared to their fixed-memory counterparts. When applied to understand the distinct bacterial phenotypic strategies observed in fluctuating nutrient environments, memory-driven cellular decision-making explains the paradoxical observation of slower growth potential in random nutrient environments when compared to constant ones. Our model predicts that this feature is universal for all fluctuating environments and memory-driven decision-making.

## Model Development

### Environmental and phenotypic dynamics

Our model consists of single cells navigating an environment characterized by type-*A* and type-*B* signals. Without loss of generality, type-*A* signals represent beneficial or nutrient-high environments, whereas type-*B* signals represent weakly beneficial or detrimental ones. Cells exist in one of two phenotypic states, correspondingly denoted by *S*_*A*_ and *S*_*B*_ based on their fitness preference for state *A* or *B*, respectively. The *S*_*A*_ state may be viewed as a fast-growing, sensitive phenotype [22]. Similarly, the *S*_*B*_ phenotype constitutively devotes additional resources to resist the undesirable effects of a nutrient-poor or overtly detrimental landscape [23]. We distinguish between *static* versus *dynamic* populations based on whether they possess the ability to switch between both states. We represent an uncertain environmental landscape *L*_*n*_ ∈ {0, 1} by a sequence of random variables indicating the presence of type-*B* signals. These signals may in general be interdependent and time inhomogeneous. For foundational understanding, we focus on environments identified by independent and identically distributed sequences *L*_*n*_ with the likelihood of type-*B* signals given by the *environmental parameter p*:

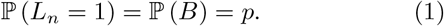

The intensity, *I*_*n*_, of each signal is in general a random quantity, which we assume to be Poisson distributed and dependent on signal type:

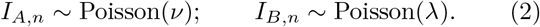

Of primary interest is the rate of benefit acquired to the cell per normalized unit time for each phenotype, given by:

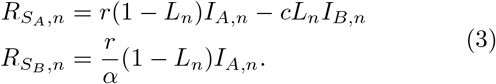

For the *S*_*A*_ phenotype, type-*A* environments generate a per-unit benefit *r*, while type-*B* signals incur a per-unit cost *c*. The *S*_*B*_ phenotype avoids this cost at the expense of less efficient navigation of type-*A* environments, parameterized by *α >* 1.

An equivalent formulation can be given in the case of strictly beneficial environments (*r* = *r*_*L*_ *< r*_*H*_ = *−c*) with *A* = ‘Low’ and *B* = ‘High’, and reversed interpretation of phenotypes *S*_*A*_ = *S*_*Low*_, *S*_*B*_ = *S*_*High*_. In this case, replacement of *α*^*−*1^ by *β >* 1 characterizes the relative efficiency of the fluctuation tolerant, slow growth *S*_*Low*_ phenotype over *S*_*High*_ in weakly beneficial environments (See SI for full details).

While the above model can represent a variety of cellular features that a system may actively optimize, including cellular signaling and phenotypic sustainment of microenvironmental features, we henceforth interpret ‘benefit’ and ‘cost’ with respect to a cell’s *growth potential*, or the rate at which cells acquire resources that are available for growth and division, which can be mapped to observable growth rates via a convex growth curve. A depiction of our phenotypic switching model is given in Fig. 1.

**Figure 1:**
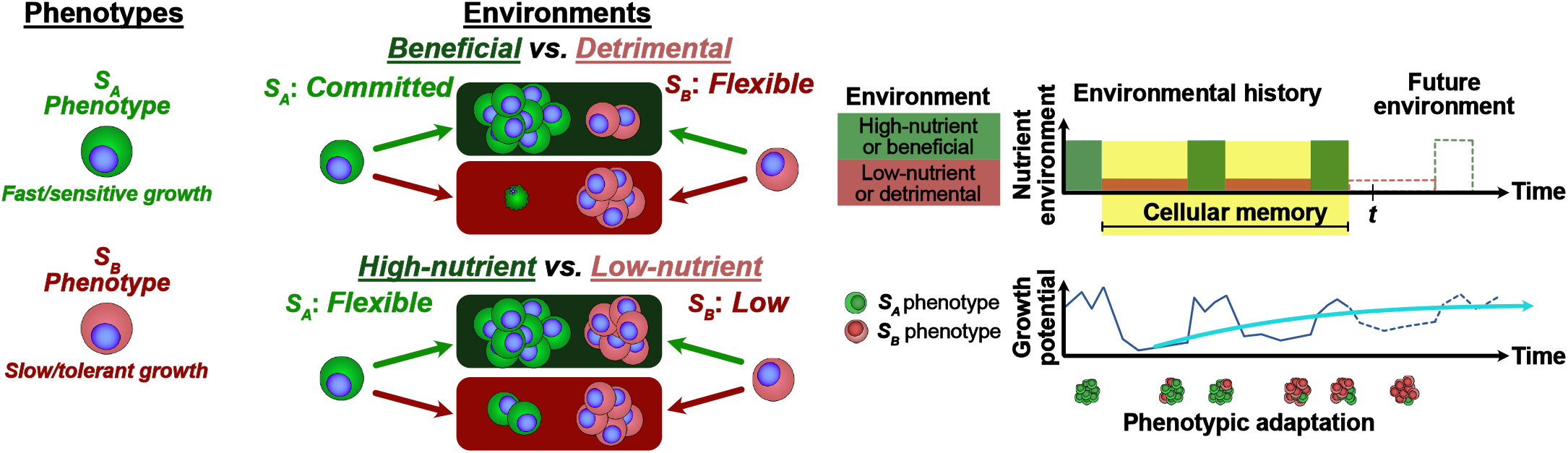
Illustration of memory-driven phenotypic switching. (A) The model consists of cells capable of stochastic transition between 2 states: *S*_*A*_ and *S*_*B*_, where each state *S*_*i*_ has an advantage in *i*-type environments. (B) The general model is applied to study cases where the environment may become hostile, or may represent low-nutrient availability. (C) Cellular decision-making occurs through the memory-driven observed occurrence rates of recently encountered environments and acts to optimally respond to future ones. Model dynamics account for the effects of cellular memory on past environmental histories in order to optimally select their phenotype while navigating uncertain future landscapes.

### Cellular memory

To navigate the *p*-fluctuating environment, the cell creates a dynamically updated estimate, *π*_*n*_, of *p* after each encountered signal. We represent iterative updating through a Bayesian inference scheme, where at time *n* the current estimate *π*_*n*_ of environment *p* may be represented by a prior distribution *f*_0_. While this framework can handle arbitrary distributions that may describe the system’s prior belief of the environmental state, we shall consider

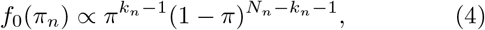

where *k*_*n*_ represents the observed number of type-*B* signals out of *N*_*n*_ recalled previous signals. In this case, the cell is tasked with identifying the most likely environment – in the form of a posterior distribution for the estimated environment *π*_*n*+1_ – given the most recent observation *L*_*n*+1_.

This distribution can be written as

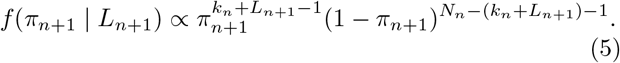

Moreover, for a uniform prior representing no bias in *π*_*n*_, the maximum likelihood estimate (MLE) of *p*, given by *π*_*n*_ = *k*_*n*_*/N*_*n*_ agrees with the maximum *a posteriori* probability (MAP), so that it is sufficient to track the number of observed type-*B* signals *k*_*n*_ for each time *n* and memory capacity *N*_*n*_. To account for the additional possibility that cells may dynamically change their memory capacity [24, 25], we consider cells possessing either a *fixed memory* capacity or an *adaptive memory* capacity as they navigate a fluctuating environment.

#### 1.1. Dynamic programming and optimal adaptation

The intriguing empirical observation of cell decision-making navigating future events based on present decisions [18, 19] is particularly well-suited for the application of stochastic optimal decision-making. Toward this end, we apply dynamic programming to describe the optimal phenotypic strategy for cells to optimize their maximal attainable expected growth potential. The optimal phenotypic policy satisfies the time-homogeneous Bellman equation [26]. This relates the *n*-period *value*– or present maximal attainable growth potential – to the value at the next period *n* + 1. Estimates of the *p* are represented by the history of type-*B* signals: *π*_*n*_ = *k*_*n*_*/N*_*n*_. The optimal program can in general be written for systems with adaptive memory (SI Sec. S5) using the value function *V*. For the fixed memory case with memory size *N* and *k*_*n*;*N*_ denoting the *N* most-recent signals at time *n*, the optimal program is given by:

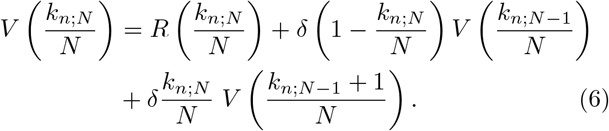

where 0 *< δ <* 1 is an exponential discount factor placing emphasis on earlier contributions to growth potential over later ones, and *R* is the expected growth reward assuming the optimal action is taken.

## Results

The following section presents the main findings of our analysis (full mathematical details are provided in the SI).

### Environmental parameters determine preferred phenotype

The mean growth potential for each phenotype is given by:

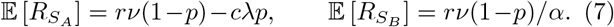

This implies that the *S*_*A*_ phenotype is preferred over the *S*_*B*_ phenotype whenever the mean benefit-to-cost ratio exceeds the odds of type-B signal arrival normalized by the inefficiency of the *S*_*B*_-phenotype:

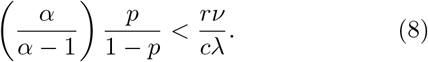

Eq. 8 equivalently describes a unique environmental landscape for which neither phenotype is preferred, given by the indifference probability *p*_*I*_ :

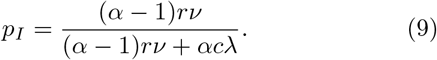

The *S*_*A*_ (resp. *S*_*B*_) phenotype is preferred in stochastic environments having *p < p*_*I*_ (resp. *p > p*_*I*_). In addition to expected dynamics, we show that there is always a variance premium incurred for the *S*_*A*_-phenotype, wherein the cost and Poisson intensity parameter terms decouple (Sec. S2.3).

### Optimal strategy and maximal total growth

The solution to the two-state problem in Eq. 6 along with its infinite memory analog can be analytically solved for arbitrary memory capacity. The per-period optimal decision *R*(*π*_*n*_) is given by selecting either a *S*_*A*_ or *S*_*B*_ phenotype based on the historical abundance of previous type-*B* signals through *π*_*n*_, and so the current growth potential is given by:

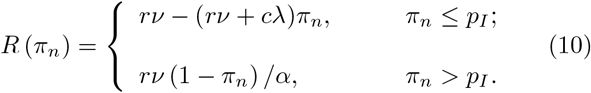

The corresponding value function, representing the maximal sum of future expected total growth potential, can be solved for in both cases (Sec. S5). In the infinite memory case, its value matches Eq. 10 normalized by 1*/*(1 *−δ*). The value function is plotted for a variety of *α* values in Fig. 2A and compared to the upper and lower limits with *α* = 1 and *α* → ∞, respectively. A similar calculation can be made for systems with finite memory (Sec. S5.1). In that case, the value function is discretized over available values of *π*, with resolution proportional to memory capacity. The finite case converges to the infinite memory one in the large memory limit (Fig. 2B). Using this updating scheme, the current growth value perceived by decision-makers in the present decreases for increased observed frequencies of the type-*B* signals, quantifying the detriment of an increasingly hostile environment. The emergent inflection point emergent for larger memories (*N*≥ 100) characterizes the relative rate of change in value for the two-environment case around the point of indifference *p*_*I*_.

**Figure 2:**
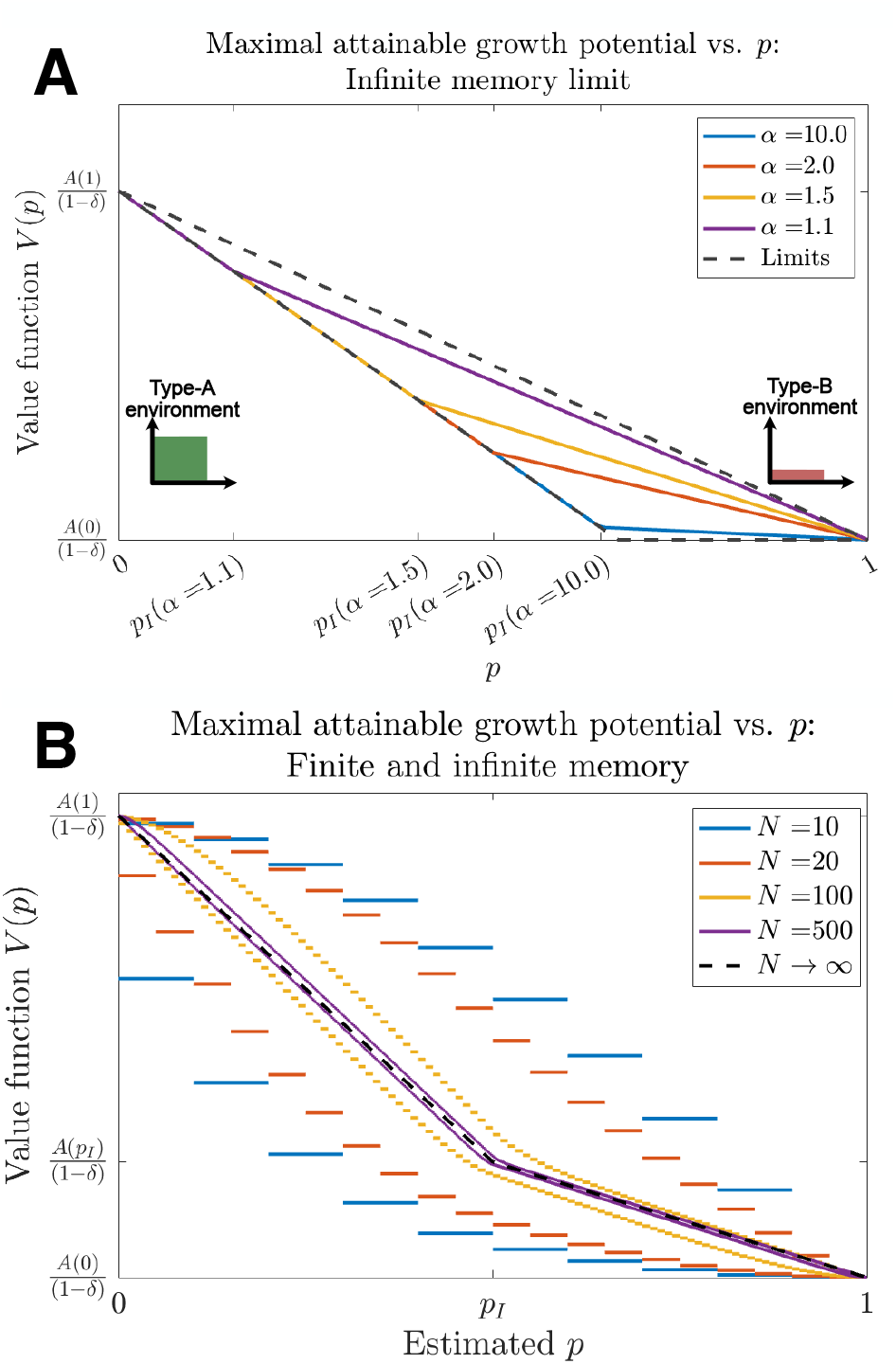
Maximal attainable growth potential in fluctuating environments. The value function for cells optimally selecting their phenotypic state based on an environmental history in a stochastic environment is given as a function of the underlying environmenttal parameter *p* for (A) systems with infinite memory for variable *α* along with upper and lower limiting behavior for *α*=1 and *α* →∞, respectively. (B) For systems with finite memory capacity, the value functions are plotted for *α*=2 over increasing memory size, along with the infinite memory limit (in all cases, *rν*=2, *cλ*=1, *δ*=0.9).

### Optimized phenotypic decision-making hedges against future environmental uncertainty

One immediate consequence of memory-driven phenotypic adaptation is that populations faced with uncertain future environments capably adapt based on estimates of the prior state. When the environment is held constant at some *p* ≠ *p*_*I*_, a corresponding fixed phenotypic state (either *S*_*A*_ or *S*_*B*_) is always preferred over phenotypic switching. Dynamic cells in unknown environments therefore experience lower growth potential relative to fixed-strategy, environmentally matched counterparts, but in doing so achieve enhanced growth potential in all environments, which contrasts with static phenotypes (Fig. 3A). This is especially relevant for detrimental type-*B* environments, wherein dynamic cells maintain positive growth despite environmental uncertainty. Thus, the benefit of phenotypic decision-making, which enables systems to survive changing environmental landscapes, is a hedge against environmental uncertainty reflected in reduced expected growth potential compared to that achieved in the static phenotype case (Fig. 3B), which converge to the predicted longterm expected behavior (Sec. S7.2). Lastly, the stochastic growth potential for static and dynamic strategies converge whenever *p* = *p*_*I*_.

**Figure 3:**
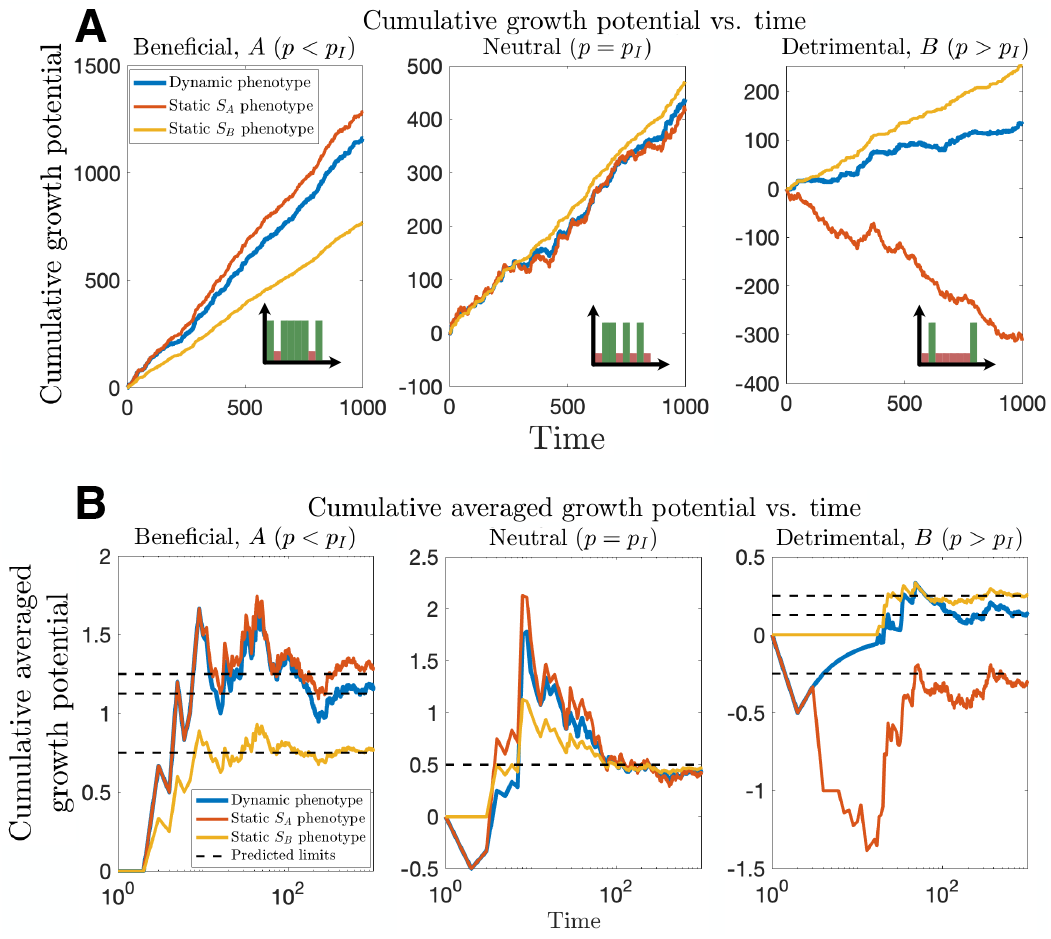
Growth in fluctuating environments. Representative stochastic trajectories of (A) cumulative growth potential and (B) averaged cumulative growth potential are depicted for cells navigating type-*A* (left), neutral (middle) and type-*B* (right) environments. Adaptive systems (blue) capable of phenotypic switching have lower growth potential than static phenotypes (red, yellow) matched to the proper environment, but outperform those phenotypes whenever the environment switched. All phenotypic strategies collapse in a neutral environment (In all cases, *rν*=2, *cλ*=1, *α*=2, *δ*=0.9 *N* =20

We next consider the role of decision-making in rich vs. deprived nutrient environments. Intriguing recent experimental evidence has described a distinct phenotype for bacterial cells navigating fluctuating high and low nutrient environments, which provides an immediate context for applying our model [21]. Motivated by this experimental setup, we for the remainder of the results focus on studying strictly beneficial high (type-*A, r* = *r*_*H*_) and low (type-*B, r* = *r*_*L*_) nutrient environments with corresponding *S*_*High*_ and *S*_*Low*_ matched phenotypes and *β* ≡*α*^*−*1^ *>* 1.

In constant environments, memory size has no influence over the preference of static or dynamic phenotypes since the environmental parameter and associated MLE always coincide (*π*_*n*_ = *p* for *p* ∈ {0, 1}). In fluctuating environments, larger memories mitigate estimation error by enhancing the resolvability of the environment relative to its indifference point, |*p −p*_*I*_| (Sec. S4.3). When switching from constant to fluctuating environments, cells with greater memory trade longer adjustment times to override memory from the prior state for greater long-term growth, resulting from reduced environmental mis-estimation in a manner that depends on the proximity of the environmental state to indifference |*p*_*I*_*− p*| (Fig. 4A, S4). Memory capacity thus determines both the rate of adaptation and long-term growth efficiency.

**Figure 4:**
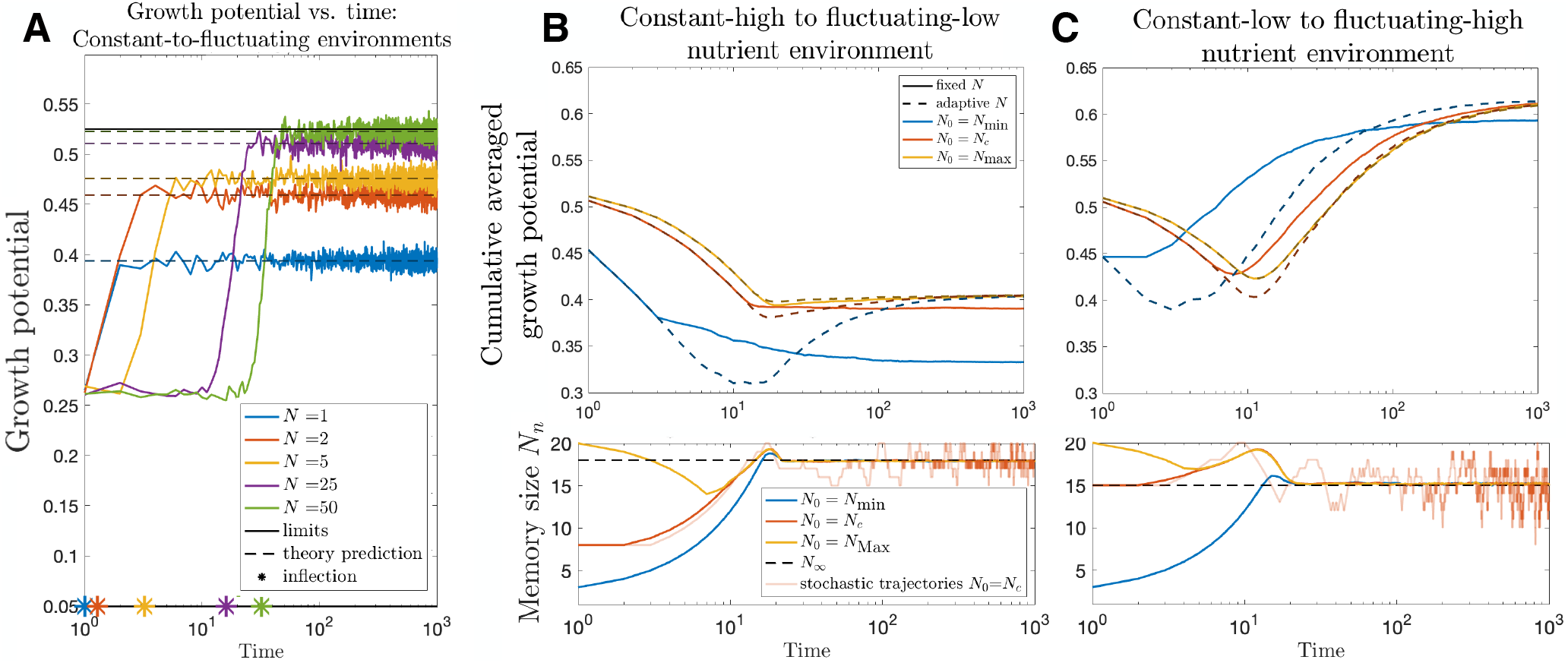
Dynamics of fixed and adaptive memory systems navigating constant-to-fluctuating nutrient environments. (A) Fixed-memory cells experiencing low-constant to high-fluctuating nutrient environments undergo phenotypic switching. The maximal growth potential and switching times both depend directly on memory capacity. Cumulative averaged growth potential and memory sizes are plotted in time for adaptive cells undergoing (B) constant-high to flutuating low and (C) constant-low to fluctuating-high nutrient environments (in all cases, *r*_*L*_ = 0.05, *r*_*H*_ = 1 giving *β*_*C*_ = 1 + *r*_*H*_ */r*_*L*_. *β* = *β*_*C*_ */*2 giving *p*_*I*_ = 0.32. For (B) and (C) *N*_min_ = 3, *N*_max_ = 20, high-fluctuating environments were given by *p* = 0.60, low-fluctuating by *p* = 0.20, and memory was updated based on deetected distance to environmental indifference *d*_*n*_ = |*π*_*n*_*− p*_*I*_|. A total of 10^4^ stochastic simulations were averaged to generate growth and memory curves. Two stochastic realizations are depicted for initial memory sizes *N*_0_ = *N*_*c*_ selected in via linear interpolation between *N*_min_ and *N*_max_ of the initial distance between *p*_*I*_ and the high- (resp. low-) constant environments, modeled by *p*_0_ = 1 (resp. 0).

### Adaptive memory balances long-term growth enhancement with short-term reduction

In our model, phenotypic dynamism enables cells to execute optimal decisions, and their accuracy is proportional to total memory capacity *N*_*n*_. Given memory’s role in cellular decision-making [15, 16], we next asked how cellular systems with dynamic memory capacities compare to their static memory counterparts when encountering fluctuating environments.

Here, memory capacity is the control variable through which the cell may tune its decision to switch phenotypes. We considered several governing principles for adaptive decision-making (Sec. S5), the one presented here is based on proximity to *p*_*I*_ as follows: Since estimation of the optimal phenotype becomes difficult for small |*p*_*I*_ *−p*|, one reasonable response for dynamical systems detecting a smaller distance *d*_*n*_ = |*π*_*n*_ *−p*_*I*_| is an increased memory capacity to more reliably estimate *p*, whereas larger *d*_*n*_ benefit from smaller memories in their ability to quickly adapt to future changes. While many functional forms can achieve this, we for simplicity considered linear interpolation between upper (*N*_*max*_) and lower (*N*_*min*_) memory limits based on proximity of *π*_*n*_ to *p*_*I*_

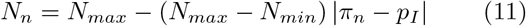

with incremental transitions so that *N*_*n*+1_ and *N*_*n*_ deviate by at most 1 unit. These dynamics describe a simple update scheme that allocates additional memory capacity when higher resolution is anticipated for resolving the environmental state. We also considered alternative schemes based on estimated variance (Sec. S6.1).

We applied this framework to compare cumulative averaged growth and memory size for systems adopting fixed versus adaptive memory strategies. Our analysis was performed for cells transitioning from a constant type-*A* environment (*p* = 0) to a type-*B*-predominant fluctuating one (*p*_*I*_ *< p <* 1) (Fig. 4B), from a constant type-*B* environment (*p* = 1) to a type-*A*-predominant fluctuating one (0 *< p < p*_*I*_) (Fig. 4C), and the corresponding fluctuating-to-constant transitions (Fig. S3). We plot the cumulative averaged growth potential to highlight both transient changes in growth during environmental switching and long-term equilibrium rates. In each case, we find that larger memory capacity improves long-term growth potential.

In constant-to-fluctuating environmental transitions, adaptive memories with lower initial memory correct more quickly than adaptive systems with large memory, demonstrating that fixed large memory capacities are not always the optimal choice (Fig. 4B,C). Fixed low-memory systems adjust more quickly, but are ultimately eclipsed by their adaptive counterparts in the long run. Moreover, adaptive systems with variable initial memory have cumulative averaged growth potentials that converge to a single predicted equilibrium state (Sec. S7.2), and a unique mean-reverting ultimate memory capacity (Sec. S6.2). Collectively, our results predict additional versatility in systems with dynamical memory capacities that is absent in static systems navigating changing environments. Moreover, initial growth reduction followed by successive enhancement with increasing time in fluctuating environments recapitulates the observed empirical behavior [21].

### Reduced growth in random environments results from inherent nutrient fluctuations and memory-dependent environmental mis-estimation

Earlier findings in Fig. 3 highlight the hedging strategy of a dynamic cellular phenotype. Namely, dynamic cells opt for lower expected growth potential relative to corresponding static phenotypes appropriately matched to a (fixed) environmental state in exchange for the ability to adapt and thus avoid large penalties in any environment of interest. Because of this, we next studied how growth under rapid fluctuation compares to the corresponding weighted average behavior between high and low states comprehensively for all random environments. An intriguing set of experiments have considered this question in bacterial decision-making. There, researchers identified a distinct phenotype under fluctuating environments with growth rates surprisingly falling short of corresponding averaged constant nutrient environments even after accounting for the convexity of growth vs. nutrient curves [21] and corresponding Jensen’s inequality.

Since our model tracks the linear underlying growth potential as a function of phenotype and environment, Jensen’s inequality holds at equality, allowing us to directly compare the growth potential of dynamic cells in fluctuating environments with the corresponding weighted average potential in constant environments (Fig. 5A). We remark that this representation of available nutrients for growth could then be mapped via one-to-one correspondence to growth rate via convex transform, but our goal is to quantify the inherent deficit in the fluctuating growth phenotype that is independent of the convexity of the growth-versus-nutrient curve. Our calculations, performed for all allowable fluctuating environments *p* and corresponding weighted-average of constant environments, recover the experimentally observed deficit of growth in the former case. Moreover, our results suggest that this deficit is universal across all feasible parameter choices (Figs. 5B,C, S6) and is greatest for near-indifferent environments *p* ≈*p*_*I*_ (Fig. 5D).

**Figure 5:**
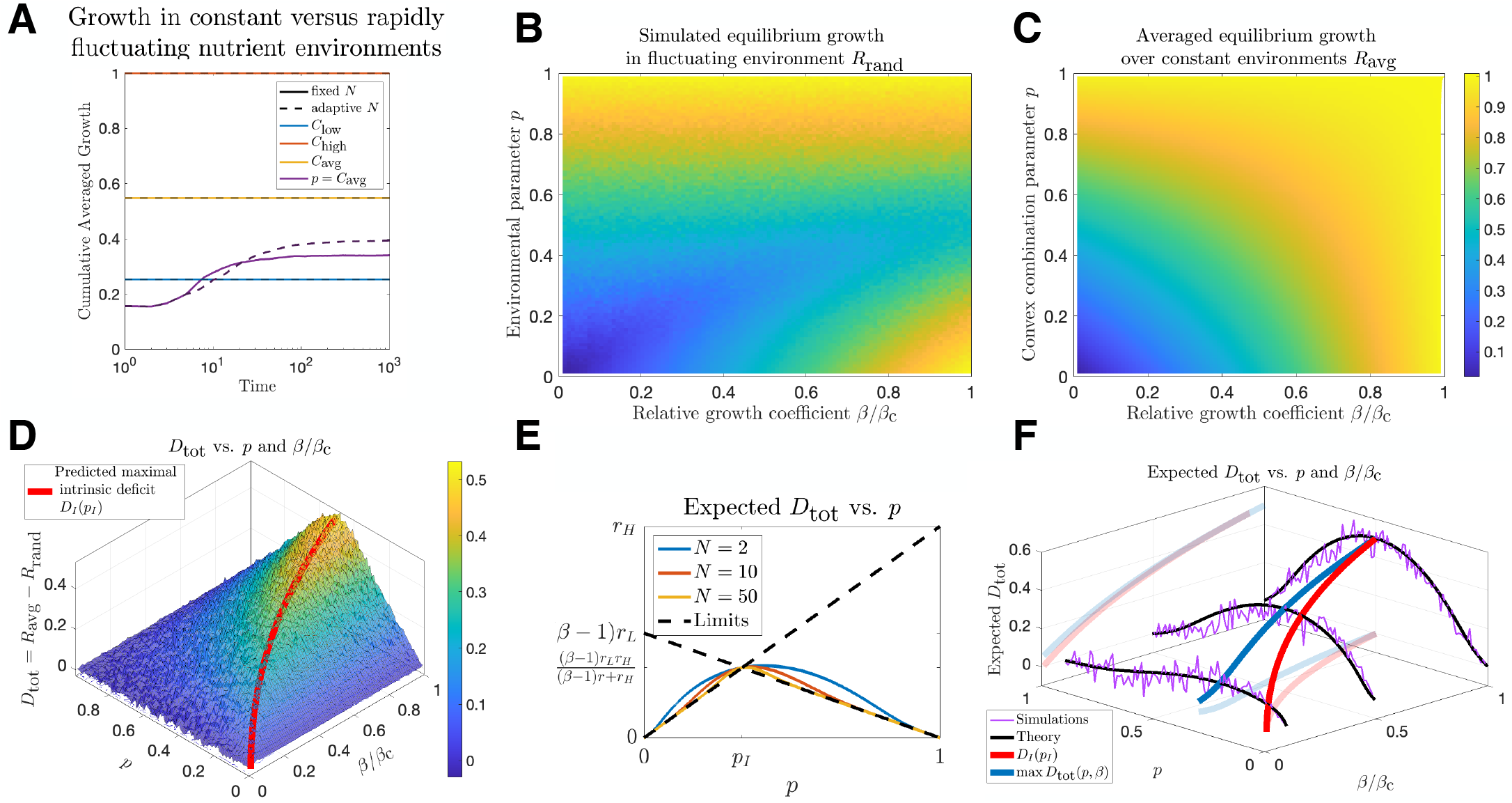
Growth deficits due to phenotypic switching in stochastic environments. *C*_avg_ describes the average environment given by the convex combination *pC*_high_ + (1 *−p*)*C*_low_ against which the rapidly fluctuating *p*-environment can be compared to. (A) The dynamics of cumulative averaged growth potential for fixed and adaptive-memory cells navigating fluctuating and comparable constant environments (*p* = 0.4, *p*_*I*_ = 0.2, *β* = *β*_Crit_*/*4). Across all environmental parameters *p* and allowable growth coefficients *β* simulated long-run growth potential for (B) *p*-random and (C) *p*-averaged constant environments reveal a universal growth deficit (D) of the *p*-fluctuating case relative to *C*_avg_, particularly pronounced in the *p* = *p*_*I*_ environment. (E, F) The total expected growth deficit of fluctuating environments consists of an intrinsic contribution maximized at *p* = *p*_*I*_ and an extrinsic contribution (illustrated for adaptive cells with limited *N* =3 memory capacity) owing to the risk of environmental phenotypic mismatch (In all cases, *r*_*L*_ = 0.01, *r*_*H*_ = 1, *β*_Crit_ = 1 + *r*_*H*_ */r*_*L*_, 10^3^ stochastic simulations were evaluated for each simulation in (A) over time and at each parameter value in (B-F)).

The total deficit can be decomposed into a sum of deterministic (*D*_*I*_) and random (*D*_*E*_) terms:

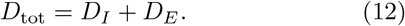

The intrinsic deficit *D*_*I*_, given by

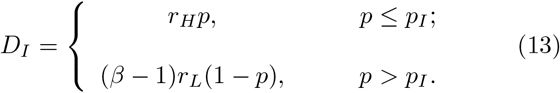

occurs when a dynamic cell under rapid environmental fluctuation correctly selects the best phenotype. The fluctuation however results in mis-match on occasion, and this effect increases as the environment approaches the state of indifference. *D*_*I*_ represents a best-case minimal deficit provided that the cell selected the correct phenotype. In the event that the phenotype is mismatched to the environment (*I*_miss_ = 1), there is an additional extrinsic deficit:

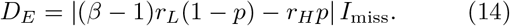

The expected value of *D*_*E*_ therefore depends on the product of its magnitude with the miss probability: The effective cost of a mismatched phenotype increases linearly away from *p*_*I*_. On the other hand the miss probability (Sec. S6) varies inversely with larger memory, is maximized at *p*_*I*_, and decreases for larger *p* deviations. This analytic description of deficit is in quantitative agreement with large-scale stochastic simulations of the process across all parameter values (Fig. 5E,F).

Collectively, our results provide a statistical justification for the observed growth deficits of fluctuating environments: Adaptive cells encountering sufficiently rapid fluctuations select their phenotypes based on the most recent past encounters. In the best case, the cell correctly identifies the most likely environment relative to *p*_*I*_, and selects the correct phenotype accordingly. In fluctuating landscapes both environmental states occur, and so growth reductions results when the anomalous environment appears despite appropriate phenotypic selection. Should the cell stochastically misestimate the environment a risk mitigated by increasing memory capacity it suffers more frequent mismatches, thus further reducing growth.

## Discussion

The dynamics underlying phenotypic cellular decision-making is central to a myriad of clinically significant phenomena, including drug resistance in infectious diseases and cancer. Analytical modeling approaches offer a direct mathematical context for studying the nature and extent of phenotypic adaptation, which can be applied to generate relevant and directly testable predictions for follow-up validation. Recent empirical findings implicate the role of phenotypic memory in adaptable systems, along with intelligent decision-making that can anticipate future events. Given this, the precise impact of prior environmental memory on improving the rate of phenotypic matching, through observable signals like growth, are of central importance.

Our predictions suggest that transient reductions in growth capacity followed by long-term optimization is an emergent defining feature of systems with dynamical memory capacity that is absent in static-memory systems. This difference presents a means by which memory-driven adaptation could be distinguished experimentally. In our model memory capacity is represented by the extent to which past encounters are remembered, which along with dynamic differences in memory size depends on the underlying molecular mechanisms giving rise to cellular memory [13, 15–17]. Their mechanistic link to phenotypic decision-making is a central question that will benefit from further experimental studies.

Here, we considered a generalized stochastic model of adaptation driven by cells choosing their optimal phenotype based on memory-driven estimation of fluctuating environments. By comparing adaptation in fluctuating environments to weighted behavior of average ones, we determined that the ability to adapt serves as a bet-hedging strategy. This causes cells undergoing rapid fluctuations to perform worse than cellular programs growing in constant conditions. Our model provides a statistical explanation for this growth deficit: The theoretically predicted extent of this deficit occurs due to the system’s existence in a stochastic environment, which inevitably experiences instances of phenotype-environmental mismatch. Moreover, our model predicts that this effect can be mitigated, but never outright eliminated, by larger memory sizes. Lastly, our results suggest the existence of fluctuating environments for which adapting cells perform most poorly, which has significant therapeutic implications for targeting adaptive threats like infectious diseases and cancer. For example, careful environmental selection based on the timescale of cellular memory could lead to growth reductions, consistent with prior observations of improved survival when choosing environmental perturbation to match the timescale of phenotypic switching [11]. This intriguing possibility will benefit from subsequent mechanistic experimental and theoretical follow-up to determine conditions under which memory-driven stochastic versus deterministic growth strategies occur. The timescales and magnitudes of changes for which cellular systems categorize their environment as either fluctuating or a new constant environment is poorly defined, and further refinement will benefit our understanding of the rules governing such transitions, which we do not address here.

In addition to recapitulating empirical findings, our results offer testable strategies for further elucidating the dynamics of memory-driven phenotypic decision-making. For example, identification of fluctuating environments giving rise to equal abundances of distinct cellular phenotypes having no significant difference in growth is one way of experimentally identifying the predicted critical indifference environment *p*_*I*_. Our results predict that cells in this environment experience large growth deficits relative to their constant-environment counterparts. Moreover, rapid environmental cycling between high and low nutrient states in a neighborhood around *p*_*I*_ could be performed to identify the environment at which the maximal deficit occurs, *p*_*max*_. Our theory suggests that deviation of *p*_*max*_ from *p*_*I*_ is an inverse readout of an adaptive system’s total memory capacity.

Since the convexity of the growth versus nutrient curve complicates quantification of phenotypic efficiency in biological systems [21], we focus on growth potential, which is a linear function of the environmental parameter. In doing so, we avoid the technicalities of invoking Jensen’s inequality when calculating phenotypic efficiency. In comparing these cases, we assumed that nutrient availability for growth in constant environments interpolates linearly between low- and high-nutrient conditions and represents a distinct phenotypic program from cells in fluctuating environments, driven in our model by experiencing rapid stochastic fluctuations.

The model presented above has several simplifying assumptions. Implicit in our modeling framework is an assumed constant energetic cost across all available memory capacities and zero net cost for transitioning. We studied memory capacities assuming instant updates between successive time-points. In reality, lags in environmental estimation and phenotypic switching may affect the optimal cellular strategy. This case can be accounted for in our framework by considering multiple arrival occurrences prior to the next phenotypic decision. Here, we assume that environmental detection and encoding into memory is perfect, when in reality this is highly dependent on the specific mechanism of memory formation in a particular system under study. Our foundational model could be extended to cases that require an account of imperfect information from the cell measuring its environment by selecting alternative functional forms for the Bayesian prior and update rules. The optimization objective we focused on was maximizing mean growth. It is possible that alternative biological settings are better represented by an alternative objective, such as variance minimization for settings requiring uniform growth, for which our approach can be generalized to describe alternative cellular stratagems.

In this work we have presented a 2-state, 2-environment model. While useful for characterizing memory-driven phenotypic transitions in some contexts [13], in others the complexity of multiple nutrient and signaling cues likely gets incorporated into the decision-making, with several allowable phenotypic states [27–29]. In these more complicated cases, the number of stable phenotypic states, along with their corresponding fitness in fluctuating landscapes, are important to evaluate. Such scenarios may utilize similar modeling to characterize optimal decision-making [30] and will benefit from data-driven model extensions to define the phenotypic response to more complex environmental signatures. At present our model does not account for durable alteration mechanisms, such as genetic mutation, that may also contribute to improved adaptation over time. Such mechanisms and their associated impact on phenotypic response is important for many evolutionary processes like cancer and is a topic of further followup investigation. Our model can be used to investigate spatiotemporal differences in environmental fluctuations inherently occurring at the micro-scale as a generator of phenotypic heterogeneity for single-cell populations and multicellular systems.

## Methods

A complete description of all mathematical details may be found in the SI Appendix.

## Acknowledgments

JTG would like to thank Kerry E. Back and Thomas J. George for their helpful discussions on stochastic dynamic programming. JTG was supported by the Cancer Prevention Research Institute of Texas (RR210080). JTG is a CPRIT Scholar in Cancer Research.

## Appendix

Supplementary information with full mathematical details are provided in the attached document.

