## Supplementary Information for "Optimal Cellular Phenotypic Adaptation in Fluctuating Environments"

### Supplementary Information: Optimal Phenotypic Adaptation in Fluctuating Environments

Jason T. George

January 17, 2023

#### S1 Overview

Here, we present the full mathematical details of our stochastic optimal cellular decision-making model. Sec. S2 describes the development of our model. Sec. S3 describes the Bayesian inference scheme utilized for stochastic environments. Sec. S4 provides the solution of the model over finite time horizons, and Sec. S5 gives the analogous solutions for systems optimizing over an infinite time horizon. Sec. S6 describes the modeling schemes used to describe adaptation of memory capacity as a function of changing environmental landscapes. Sec. S7 discusses the application of our model to fluctuating nutrient-high and low environments. Sec. S8 contains a list of supplementary figures.

### S2 Model development

We consider the following stochastic model: A cell receives an external signal at some fixed rate per unit time. The signal is either beneficial (type-$A$) or is detrimental (type-$B$). The magnitude of a beneficial signal may in general be Poisson-distributed with parameter $\nu$ scaled by a per-unit reward $r$. Similarly, the cost of each detrimental signal is Poisson-distributed with parameter $\lambda$ scaled by a unit cost $c$. Type-$B$ signals arrive independently with probability $p$, and we refer to this as the *environmental parameter*.

We consider the scenario wherein adaptive populations navigate fluctuating environments by selecting their phenotype based on a memory of past environmental occurrences. We distinguish this decision-making state from cells experiencing constant (non-random) environments in line with empirical observation [1]. Between the arrival of each signal, a cell may choose to exist in one of two states. In the committed phenotype, $S_A$, a cell can efficiently process the signal it receives over $\Delta t$ time. In the flexible phenotype, denoted by $S_B$, a cell is resistant to an alternative signal (which may result from a phenotype capable of processing it, for example). We will henceforth refer to cells adopting the $S_B$ flexible phenotype as *surveillance*. The ability to process both types of signal of course requires a diversion of resources otherwise freely available in the committed phenotype, and so we assume that it takes $\alpha\Delta t$ time to process the signal, with $\alpha > 1$ (or, equivalently, that less resource is available per unit time while flexibly adapting).

We start by considering the conditions under which the committed $S_A$ phenotype is preferred or even feasible. Ultimately, we will be interested in studying the optimal strategy that the cell may take if the underlying frequency of alternative signals $p$ is unknown to the cell. We let $L_n$ denote the sequence of independent, identically distributed (IID) indicator random variables for the presence of a type-$B$ signal at period $n$. Each $L_n$ is distributed Bernoulli($p$). We denote by $I_{A,n}\sim$Poisson($\nu$) the benefit of a beneficial type-$A$ signal, and by $I_{B,n}\sim$ Poisson($\lambda$) the cost of a type-$B$ signal.

#### S2.1 Expected growth potential

If maximizing the total accrued beneficial signal were all that mattered, then surveillance would be preferred at all times. However, given that surveillance takes additional processing time (or a reduced amount of resources is freely available to the cell for the same fixed time interval) relative to the non-surveillance option, we consider the rate of signal accrual. We study the growth potential, which corresponds to the extent of resources available for growth per normalized unit time interval $\Delta t$. Let $R_{S_B,n}$ (resp. $R_{S_A,n}$) be the growth potential under surveillance (resp. no surveillance). Then:

$$R_{S_A,n} = r\big(1 - L_n\big)I_{A,n} - cL_nI_{B,n}; \qquad R_{S_B,n} = \frac{r}{\alpha}\Big(1 - L_n\Big)I_A. \tag{S1}$$

The expected growth potential experienced by the $S_A$ phenotype (under no surveillance) is

$$\mathbb{E}\left[R_{S_A,n}\right] = r\big(1 - \mathbb{E}\left[L_n\right]\big)\mathbb{E}\left[I_{A,n}\right] - c\mathbb{E}\left[L_n\right]\mathbb{E}\left[I_{B,n}\right] = r\nu(1-p) - c\lambda p. \tag{S2}$$

Viability of no surveillance occurs whenever Eq. S2 is positive and requires that the odds of a type-$B$ signal be bounded above:

$$\frac{p}{1-p} < \frac{r\nu}{c\lambda}. \tag{S3}$$

We may alternatively derive the viability probability, or maximal environmental value for which the $S_A$ phenotype is viable:

$$p_{viable} = \frac{r\nu}{r\nu + c\lambda}. \tag{S4}$$

The expected growth potential experienced by the $S_B$ phenotype (under surveillance) is

$$\mathbb{E}\left[R_{S_B,n}\right] = r\nu(1-p)/\alpha. \tag{S5}$$

From this, it is clear that the $S_A$ phenotype is preferred over the $S_B$ phenotype if

$$r\nu(1-p) - c\lambda p > r\nu(1-p)/\alpha \qquad \Leftrightarrow \qquad \left(\frac{\alpha-1}{\alpha}\right)\frac{r\nu}{c\lambda} > \frac{p}{1-p}. \tag{S6}$$

Eq. S6 states that the odds of a type-$B$ signal must be further reduced in a manner reflected in the relative inefficiency of surveillance $\alpha$. From Eq. S6 we may identify the *critical indifference probability*, $p_I$ for which the system has no phenotypic preference:

$$p_I = \frac{(\alpha-1)r\nu}{(\alpha-1)r\nu + \alpha c\lambda}. \tag{S7}$$

#### S2.2 Beneficial signals

The above analysis proceeds identically if instead of incurring a cost in the alternative state, the cell receives a low-growth signal $r_L$, and in the beneficial state it receives a high-growth signal $r_H$, with $r_L < r_H$. In this case, the cell may similarly choose from one of two phenotypes: either a flexible high-environment phenotype, $S_{High}$ that accepts both signals, or a $S_{Low}$ phenotype economically specialized to handle growth in the $r_L$ environment with an efficiency coefficient of $\beta > 1$. For these cases the growth rates per unit time become

$$R_{S_F,n} = r_L\big(1-L_n\big)I_{L,n} + r_H H_n I_{H,n}, \qquad R_{S_L,n} = \beta r_L\big(1 - H_n I_{L,n}\big). \tag{S8}$$

We note that these are none other than Eqs. S1 with the new flexible phenotype ($S_F$)replacing the original committed ($S_A$) one, and the new economical low-growth phenotype ($S_L$) replacing the original flexible one ($S_B$), $H_n$ indicating the $H$ environment, and $I_{H,n}$ (resp. $I_{L,n}$) indicating the magnitude of a high (resp. low) growth signal. Here, Eq. S7 and the subsequent analysis all hold. We will consider this case in more detail in Secs. S6,S7. Until then, we consider positive incurred cost in the alternative environment.

#### S2.3 Growth potential variation

The variance in the growth potential for the $S_A$ phenotype,

$$\mathbb{V}\text{ar}\left(R_{S_A,n}\right) = r^2\mathbb{V}\text{ar}\left((1-L_n)I_{A,n}\right) + c^2\mathbb{V}\text{ar}\left(L_n I_{B,n}\right) - 2\mathbb{C}\text{ov}\Big(r(1-L_n)I_{A,n},\ cL_n I_{B,n}\Big), \tag{S9}$$

can be calculated from the following first and second moments:

$$\mathbb{E}\left[(1-L_n)^2 I_{A,n}^2\right] = (\nu+\nu^2)(1-p), \tag{S10}$$

$$\mathbb{E}\left[(1-L_n)L_{A,n}\right]^2 = \nu^2(1-p), \tag{S11}$$

$$\mathbb{E}\left[L_n^2 I_{B,n}^2\right] = (\lambda+\lambda^2)p, \ \text{ and} \tag{S12}$$

$$\mathbb{E}\left[L_n I_{B,n}\right]^2 = (\lambda p)^2. \tag{S13}$$

This ultimately yields

$$\mathbb{V}\text{ar}\left((1-L_n)I_{A,n}\right) = \nu(1-p) + \nu^2 p(1-p) \tag{S14}$$

$$\mathbb{V}\text{ar}\left(L_n I_{B,n}\right) = \lambda p + \lambda^2 p(1-p), \tag{S15}$$

Thus the variance under no surveillance becomes:

$$\mathbb{V}\text{ar}\left(R_{S_A,n}\right) = r^2\nu(1-p) + c^2\lambda p + (r\nu + c\lambda)^2 p(1-p). \tag{S16}$$

Similarly, we find that for the variance in growth potential under surveillance ($S_B$),

$$\mathbb{V}\text{ar}\left(R_{S_B,n}\right) = \left(\frac{r}{\alpha}\right)^2\mathbb{V}\text{ar}\left((1-L_n)I_{A,n}\right) = \left(\frac{r}{\alpha}\right)^2\nu(1-p) + \left(\frac{r\nu}{\alpha}\right)^2 p(1-p) < \mathbb{V}\text{ar}\left(R_{S_A,n}\right). \tag{S17}$$

Therefore, there is always a variance premium for no surveillance:

$$\begin{aligned}\mathbb{V}\text{ar}\left(R_{S_A,n}\right) &= \alpha^2\mathbb{V}\text{ar}\left(R_{S_B,n}\right) + c^2\lambda p + \left[(c\lambda)^2 + 2r\nu c\lambda\right]p(1-p)\\ &= \alpha^2\mathbb{V}\text{ar}\left(R_{S_B,n}\right) + c\lambda p\left[c + (c\lambda + 2r\nu)(1-p)\right].\end{aligned} \tag{S18}$$

#### S3 Bayesian inference update

The cellular decision-making process may be represented in general using a Bayesian inference-based scheme as follows: With the goal of optimally estimating the unknown environmental parameter $p \in [0,1]$ by $\pi$, the cell may initially possess a prior belief about the environment, which is represented by a prior distribution over $[0,1]$. The conjugate prior for a sequence of Bernoulli observations is the most useful representation for this distribution, which is a beta distribution:

$$f_0(\pi) \propto \pi^{a-1}(1-\pi)^{b-1}, \;\; 0 \leq \pi \leq 1. \tag{S19}$$

The functional form of Eq. S19 is also a natural one for incorporating knowledge of past environmental landscapes, for if the prior history contains $\ell$ out of $m$ alternative signals, then the corresponding prior $f_0(\pi) \propto \pi^{\ell}(1-\pi)^{m-\ell}$ is represented by letting $a \equiv \ell + 1$ and $b \equiv m - \ell + 1$. Moreover, a fluctuating environment can be written as a sum $Y_n = X_1 + X_2 + \cdots + X_n$ of $n$ independent $X_i \sim \text{Bernoulli}(\pi)$ random variables, each indicating the presence or absence of an alternative signal at period $i$. The distribution of $Y_n$ given $\pi$ is

$$f(Y_n = k \mid \pi) = \pi^k(1-\pi)^{n-k}. \tag{S20}$$

By Bayes' Theorem, the posterior distribution for $\pi$ given the observed $Y$ is

$$f(\pi \mid Y_n) = \frac{f(Y_n \mid \pi) f_0(\pi)}{f(Y_n)} \tag{S21}$$

$$\propto \pi^{a-1+k}(1-\pi)^{b-1+n-k}. \tag{S22}$$

We remark that the absence of any historical belief about $p$ may be represented in Eq. S19 with $a = b = 1$, for which $f_0$ is uniform. Moreover, for a uniform prior $f_0 = 1$, the maximum likelihood point-estimate (MLE) of $p$, given by $\pi = k/n$, agrees with the maximum a posteriori probability (MAP). In summary, in characterizing dynamic cellular decision-making with updates, it suffices to track the number of alternative and total observed signals. Simulated stochastic trajectories for growth potential and simulated population size for static and dynamic phenotypes in beneficial, neutral, and detrimental environments are plotted in Figs. 3, S1.

### S4 Infinite memory over a finite time horizon

The environmental parameter $p$ is unknown to the cell, which observes and processes arrivals as they appear. Between each arrival, the cell must determine the most suitable phenotype for the next period. We assume that the cell aims to maximize its expected growth rate while receiving a total of $N$ signals and perceives the environment based on this history. Here, infinite memory capacity means that all previous environmental exposures are recalled by the cell each time a new decision is made.

#### S4.1 Stochastic dynamic programming

Our goal is to find the maximal attainable *value*, or sum of time-discounted expected future growth potentials assuming the optimal strategy is taken, and corresponding optimal decision policy as a cell gains additional information about a fluctuating environment. Such approaches have been employed previously to describe optimal strategies in cancer progression [2]. We consider state space $\mathcal{S} = \mathbb{Q} \cap [0, 1]$, where state $\pi$ denotes the current estimate $p$. We denote by $V_n(\pi)$ the maximal attainable value after processing $n$ total signals when in state $\pi$. The current state may be represented by the maximum likelihood point-estimate of $p$ (unknown to the population), given by $\pi = k/n$, where $k$ is the current number encountered type-$B$ signals in a recalled history of $n$ total signals, and $k \leq n \leq N$. The action set is $\mathcal{A} = \{0, 1\}$, where action $a = 1$ (resp. $a = 0$) represents the cell residing in the $S_B$ state with surveillance (resp. the $S_A$ state with no surveillance). For this system, the Bellman equation may be written as

$$V_n\left(\frac{k}{n}\right) = \max_{a \in \mathcal{A}} \left\{ R\left(\frac{k}{n}, a\right) + \delta \left[ \left(\frac{k}{n}\right) V_{n+1}\left(\frac{k+1}{n+1}\right) + \left(1 - \frac{k}{n}\right) V_{n+1}\left(\frac{k}{n+1}\right) \right] \right\}, \tag{S23}$$

where $R(\pi, a)$ denotes the expected growth potential in state $\pi$ when taking action $a$, and $0 < \delta \leq 1$ is a discount to (optionally) prioritize present growth over future growth. In this particular problem, the optimal decision and reward decouple from the future value giving a time-additive recurrence relation:

$$V_n\left(\frac{k}{n}\right) = \max_{a \in \mathcal{A}} \left\{ R\left(\frac{k}{n}, a\right) \right\} + \delta \left[ \left(\frac{k}{n}\right) V_{n+1}\left(\frac{k+1}{n+1}\right) + \left(1 - \frac{k}{n}\right) V_{n+1}\left(\frac{k}{n+1}\right) \right]. \tag{S24}$$

#### S4.2 Optimal policy

From Eqs. S2,S5,S7, we have that

$$\max_{a \in \mathcal{A}} \left\{ R\left(\frac{k}{n}, a\right) \right\} \equiv A\left(\frac{k}{n}\right) = \begin{cases} A_{S_B}\left(\frac{k}{n}\right) = \frac{r\nu}{\alpha}\left(1 - \frac{k}{n}\right), & \frac{k}{n} > p_I; \\ A_{S_A}\left(\frac{k}{n}\right) = r\nu\left(1 - \frac{k}{n}\right) - c\lambda\left(\frac{k}{n}\right), & \frac{k}{n} \leq p_I. \end{cases} \tag{S25}$$

The optimal policy is in this way determined at each period based on the history of encountered alternative signals. The optimality equation can thus be written as

$$V_n\left(\frac{k}{n}\right) = A\left(\frac{k}{n}\right) + \delta \left[ \left(\frac{k}{n}\right) V_{n+1}\left(\frac{k+1}{n+1}\right) + \left(1 - \frac{k}{n}\right) V_{n+1}\left(\frac{k}{n+1}\right) \right], \tag{S26}$$

subject to the boundary condition:

$$V_N\left(\frac{k}{N}\right) = A\left(\frac{k}{N}\right), \qquad k \in \{0, 1, ..., N\}. \tag{S27}$$

Using backward induction, we find that

$$V_{N-1}\left(\frac{k-1}{N-1}\right) = A\left(\frac{k-1}{N-1}\right) + \delta\frac{N-k}{N-1} A\left(\frac{k-1}{N}\right) + \delta\frac{k-1}{N-1} A\left(\frac{k}{N}\right), \tag{S28}$$

and similarly

$$V_{N-2}\left(\frac{k-2}{N-2}\right) = A\left(\frac{k-2}{N-2}\right) + \frac{\delta}{N-2}\left[(N-k)A\left(\frac{k-2}{N-1}\right) + (k-2)A\left(\frac{k-1}{N-1}\right)\right]$$
$$+ \frac{\delta^2}{(N-2)(N-1)}\left[(N-k)(N-k+1)A\left(\frac{k-2}{N}\right)\right.$$
$$\left. + 2(N-k)(k-2)A\left(\frac{k-1}{N}\right) + (k-2)(k-1)A\left(\frac{k}{N}\right)\right]. \tag{S29}$$

Continuing, we ultimately obtain

$$V_{N-m}\left(\frac{k-m}{N-m}\right) = \sum_{i=0}^{m}\delta^i \frac{(N-m-1)!}{(N-m-1+i)!}\sum_{j=0}^{i}\binom{i}{j}\frac{(N-k-1+i-j)!}{(N-k-1)!}\frac{(k-m-1+j)!}{(k-m-1)!}A\left(\frac{k-m+j}{N-m+i}\right). \tag{S30}$$

#### S4.3 Erroneous decision likelihood

For environments driven by a constant $p$, the Strong Law of Large Numbers guarantees the convergence $\pi \overset{a.s.}{\rightarrow} p$. The proximity of the underlying environmental parameter $p$ to the indifference probability $p_I$ is thus an important quantity affecting the likelihood of stochastically generated erroneous decision-making. Since the number of type-$B$ signals $k$ observed after a total of $N$ is distributed Binomial($N$,$p$), the state variable $\pi = k/N$ is also binomial with

$$\mathbb{E}\left[\frac{k}{N}\right] = p, \qquad \mathbb{V}\text{ar}\left(\frac{k}{N}\right) = \frac{p(1-p)}{N}. \tag{S31}$$

The distance between $p$ and $p_I$ is given by

$$|p_I - p| = \left|\frac{(\alpha-1)r\nu}{(\alpha-1)r\nu + \alpha c\lambda} - p\right| = \frac{|(\alpha-1)r\nu(1-p) - \alpha c\lambda p|}{(\alpha-1)r\nu + \alpha c\lambda}. \tag{S32}$$

By Chebyshev's Inequality, we may obtain an upper bound on the probability that an erroneous decision is made after $N$ observations:

$$\mathbb{P}\left(\left|\frac{k}{N} - p\right| > |p_I - p|\right) \leq \frac{p(1-p)}{N}\left(\frac{(\alpha-1)r\nu - \alpha c\lambda}{(\alpha-1)r\nu(1-p) - \alpha c\lambda p}\right)^2. \tag{S33}$$

### S5 Infinite time horizon optimization

This case considers a cellular agent navigating a fluctuating environment over many periods, which is relevant in the case where a long-lived cell must make many decisions prior to division or death. We can consider both the case where the cell has a fixed prior memory capacity as well as the theoretical limit of infinite memory. For this case, we study the time-homogeneous Bellman equation. We denote by $k_{n;N_n}$ the frequency of type-$B$ signals observed out of the most recent $N_n$ previous signals at time $n$. The Bellman equation is

$$V\left(\frac{k_{n;N_n}}{N_n}\right) = \max_{a\in\mathcal{A}}\left\{R\left(\frac{k_{n;N_n}}{N_n}, a\right)\right\} + \delta\left[\left(1-\frac{k_{n;N_n}}{N_n}\right)V\left(\frac{k_{n;N_{n+1}-1}}{N_{n+1}}\right) + \frac{k_{n,N_n}}{N_n}V\left(\frac{k_{n;N_{n+1}-1}+1}{N_{n+1}}\right)\right] \tag{S34}$$

with $0 \leq k_n \leq N_n \leq n$. Here, strict discounting $(0 < \delta < 1)$ is required for a non-degenerate solution, since Eq. S34 evaluated at $k_{n;N_n} = 0$ yields

$$V(0) = \max_{a\in\mathcal{A}} R(0, a) + \delta V(0). \tag{S35}$$

#### S5.1 Finite memory

Eq. S34 describes the general case where memory capacity may be chosen arbitrarily at each time so long as $N_n \leq n$. We henceforth consider a system having a finite and fixed memory capacity able to recall at most $N$ of the $n$ prior states, so that

$$N_n = \begin{cases} n, & n < N; \\ N, & n \geq N. \end{cases} \tag{S36}$$

We consider some time $n \geq N$ so that the system is operating at memory capacity. Eq. S34 can be written more succinctly in this case as

$$V\left(\frac{k_{n;N}}{N}\right) = R\left(\frac{k_{n;N}}{N}\right) + \delta\left(1-\frac{k_{n;N}}{N}\right)V\left(\frac{k_{n;N-1}}{N}\right) + \delta\frac{k_{n;N}}{N}V\left(\frac{k_{n;N-1}+1}{N}\right).$$

We let $K_i$ be a 0-1 random variable indicating if a type-$B$ signal occurred at period $i$, and similarly $K_{i,j}$ be the sum of such random variables from periods $i$ to $j$. If we assume an infinite horizon optimization with $N$ recalled prior memories, Eq. S37 becomes

$$\begin{aligned} V\left(\frac{K_{n-N+1,n}}{N}\right) = A\left(\frac{K_{n-N+1,n}}{N}\right) + \delta\Bigg[&\left(1-\frac{K_{n-N+1,n}}{N}\right)V\left(\frac{K_{n-N+2,n}+0}{N}\right) \\ &+\left(\frac{K_{n-N+1,n}}{N}\right)V\left(\frac{K_{n-N+2,n}+1}{N}\right)\Bigg], \end{aligned} \tag{S37}$$

Recognizing that $K_{n-N+1,n} = K_{n-N+1} + K_{n-N+2,n}$, we obtain two recursive relations for the value function by conditioning on the value of $K_{n-N+1}$ (the oldest memory that is replaced by the next outcome).

**Boundary conditions:** If $K_{n-N+1,n} = 0$, then Eq. S37 reduces to the lower boundary condition:

$$V(0) = A(0) + \delta V(0) \qquad \Rightarrow \qquad V(0) = (1-\delta)^{-1}A(0). \tag{S38}$$

On the other hand, $K_{n-N+1,n} = N$, together with Eq. S37 gives the upper boundary condition:

$$V(1) = A(1) + \delta V(1) \qquad \Rightarrow \qquad V(1) = (1-\delta)^{-1}A(1). \tag{S39}$$

**Case I:** $K_{n-N+1} = 0$. In this case, Eq. S37 becomes

$$V\left(\frac{K_{n-N+2,n}}{N}\right) = A\left(\frac{K_{n-N+2,n}}{N}\right) + \delta\Bigg[\left(1 - \frac{K_{n-N+2,n}}{N}\right)V\left(\frac{K_{n-N+2,n}+0}{N}\right) + \left(\frac{K_{n-N+2,n}}{N}\right)V\left(\frac{K_{n-N+2,n}+1}{N}\right)\Bigg]. \tag{S40}$$

Conditioning on $K_{n-N+2,n} = k$, $1 \leq k \leq N-1$ gives

$$V\left(\frac{k}{N}\right) = A\left(\frac{k}{N}\right) + \delta\left(1 - \frac{k}{N}\right)V\left(\frac{k}{N}\right) + \frac{\delta k}{N}V\left(\frac{k+1}{N}\right), \tag{S41}$$

from which we may derive a backward recursive relation:

$$V\left(\frac{k}{N}\right) = \frac{1}{N + (N-k)\delta}\left[NA\left(\frac{k}{N}\right) + \delta k V\left(\frac{k+1}{N}\right)\right]. \tag{S42}$$

**Case II:** $K_{n-N+1} = 1$. Again conditioning on $K_{n-N+2,n} = k$, $1 \leq k \leq N-1$, Eq. S37 becomes

$$V\left(\frac{k+1}{N}\right) = A\left(\frac{k+1}{N}\right) + \delta\left(1 - \frac{k+1}{N}\right)V\left(\frac{k}{N}\right) + \frac{\delta(k+1)}{N}V\left(\frac{k+1}{N}\right), \tag{S43}$$

which yields a forward recursive relation:

$$V\left(\frac{k+1}{N}\right) = \frac{1}{N - (k+1)\delta}\left[NA\left(\frac{k+1}{N}\right) + \delta(N-k-1)V\left(\frac{k}{N}\right)\right]. \tag{S44}$$

Together, the above cases permit an exact solution for the value function. If $K_{n-N+1} = 1$, then

$$V\left(\frac{k}{N}\right) = \left(\frac{N}{N-k\delta}\right)\left[A\left(\frac{k}{N}\right) + \sum_{j=1}^{k-1}\delta^j A\left(\frac{k-j}{N}\right)\prod_{i=0}^{j-1}\frac{(N-k+i)}{N-(k-i-1)\delta}\right] + A(0)\left(\frac{\delta^k}{1-\delta}\right)\prod_{j=1}^{k}\left(\frac{N-j}{N-j\delta}\right). \tag{S45}$$

If $K_{n-N} = 0$, then

$$V\left(\frac{N-k}{N}\right) = \left(\frac{N}{N-k\delta}\right)\left[A\left(\frac{N-k}{N}\right) + \sum_{j=1}^{k-1}\delta^j A\left(\frac{N-(k-j)}{N}\right)\prod_{i=0}^{j-1}\frac{(N-k+i)}{N-(k-i-1)\delta}\right] + A(1)\left(\frac{\delta^k}{1-\delta}\right)\prod_{j=1}^{k}\left(\frac{N-j}{N-j\delta}\right). \tag{S46}$$

(S47)

Collectively, together with the fact that $A(1) = 0$, we ultimately obtain

$$
V\left(\frac{k}{N}\right)=\begin{cases}\left(\dfrac{N}{N-k\delta}\right)\left[A\left(\dfrac{k}{N}\right)+\displaystyle\sum_{i=1}^{k-1}\delta^i A\left(\dfrac{k-i}{N}\right)\prod_{j=0}^{i-1}\dfrac{N-k+j}{N-(k-j-1)\delta}\right] & \\ +\left(\dfrac{\delta^k}{1-\delta}\right)\left(\displaystyle\prod_{j=1}^{k}\dfrac{N-j}{N-j\delta}\right)\left(\dfrac{r\nu}{1-\delta}\right), & K_{n-N+1}=1; \\ & \\ \left(\dfrac{N}{N(1-\delta)+k\delta}\right)\left[A\left(\dfrac{k}{N}\right)+\displaystyle\sum_{i=1}^{N-k-1}\delta^i A\left(\dfrac{k+i}{N}\right)\prod_{j=0}^{i-1}\dfrac{(k+j)}{N(1-\delta)+(k+j+1)\delta}\right], & K_{n-N+1}=0.\end{cases} \tag{S48}
$$

##### S5.1.1 Special Case: Single-period memory

In the $N = 1$ case, the value function only depends on the prior outcome $K_{n-1}$ so that

$$
V\left(K_{n-1}\right)=A\left(K_{n-1}\right)+\delta\left(1-K_{n-1}\right)V(0)+\delta K_{n-1}V(1). \tag{S49}
$$

Here, conditioning on $K_{n-1} = 0$ and $K_{n-1} = 1$ gives the lower and upper boundary conditions of Eqs. S38 and S39, respectively. Thus,

$$
V\left(K_{n-1}\right)=(1-K_{n-1})r\nu(1-\delta)^{-1} \tag{S50}
$$

This is an extreme case where the cell's estimate $\pi$ of environmental parameter $p$ is maximally coarse so that $\pi \in \{0, 1\}$. This guarantees incorrect inference for all but trivial values of $p$. In the other extreme case, the domain of the value function $\pi$ may take any rational value on the interval $[0, 1]$. This occurs in the limit of an infinite memory process and is considered below.

#### S5.2 Infinite memory

If the system may contain a memory of the entire prior history of $M$ observations, then the stationary Bellman equation is of the form:

$$
V\left(\frac{k}{N}\right)=A\left(\frac{k}{N}\right)+\delta\left(1-\frac{k}{N}\right)V\left(\frac{k}{N+1}\right)+\delta\left(\frac{k}{N}\right)V\left(\frac{k+1}{N+1}\right). \tag{S51}
$$

Now, clearly, if $k = 0$, then Eq. S51 gives

$$
V(0)=A(0)+\delta V(0) \qquad \Rightarrow\ V(0)=r\nu(1-\delta)^{-1} \tag{S52}
$$

Similarly, for $k = N$, Eq. S51 yields

$$
V(1)=A(1)+\delta V(1) \qquad \Rightarrow\ V(1)=A(1)(1-\delta)^{-1}=0 \tag{S53}
$$

Lastly, we note that at the point of indifference,

$$
x_I=\frac{(\alpha-1)r\nu}{(\alpha-1)r\nu+\alpha c\lambda}, \tag{S54}
$$

we have

$$
A(x_I)=\frac{r\nu c\lambda}{(\alpha-1)r\nu+\alpha c\lambda}. \tag{S55}
$$

Intuitively, we may therefore guess that the value function is defined by the line segments connecting $\big(0, r\nu(1-\delta)^{-1}\big)$ to $\big(x_I, A(x_I)(1-\delta)^{-1}\big)$, and $\big(x_I, A(x_I)(1-\delta)^{-1}\big)$ to $\big(1, 0\big)$, which motivates the following result:

**Proposition 1.** *The value function in the infinite memory model is given by*

$$V(x) = \begin{cases} \frac{r\nu}{1-\delta}\left(1 - \frac{r\nu+c\lambda}{r\nu}x\right), & x \leq x_I; \\ \\ \frac{r\nu}{\alpha(1-\delta)}\left(1 - x\right), & x > x_I. \end{cases} \tag{S56}$$

*Proof.* We show that Eq. S51 holds for the following two cases, with $x = k/N$:

**Case I: $\boldsymbol{x \leq x_I}$.** The right-hand side of Eq. S51 becomes

$$r\nu - (r\nu + c\lambda)\frac{k}{N} + \delta\left(1 - \frac{k}{N}\right)\frac{r\nu}{1-\delta}\left[1 - \left(\frac{r\nu + c\lambda}{r\nu}\right)\frac{k}{N+1}\right] + \delta\frac{k}{N}\frac{r\nu}{1-\delta}\left[1 - \left(\frac{r\nu + c\lambda}{r\nu}\right)\frac{k+1}{N+1}\right]. \tag{S57}$$

factoring and rearrangement gives

$$\frac{r\nu}{1-\delta}\left[(1-\delta)\left(1 - \frac{r\nu + c\lambda}{r\nu}\frac{k}{N}\right) + \delta - \delta\left(\frac{r\nu + c\lambda}{r\nu}\right)\left(\frac{k}{N+1} + \frac{k}{N(N+1)}\right)\right]. \tag{S58}$$

Recognizing that

$$\frac{k}{N+1} + \frac{k}{N(N+1)} = \frac{k}{N} \tag{S59}$$

together with further simplification ultimately gives

$$\frac{r\nu}{1-\delta}\left[1 - \left(\frac{r\nu + c\lambda}{r\nu}\right)\frac{k}{N}\right], \tag{S60}$$

which is none other than $V(x)$, for $x \leq x_I$.

**Case II: $\boldsymbol{x > x_I}$.** A similar strategy in this case gives, for the right-hand side of Eq. S51

$$\frac{r\nu}{\alpha}\left(1 - \frac{k}{N}\right) + \delta\left(1 - \frac{k}{N}\right)\frac{r\nu}{\alpha(1-\delta)}\left(1 - \frac{k}{N+1}\right) + \delta\frac{k}{N}\frac{r\nu}{\alpha(1-\delta)}\left(1 - \frac{k+1}{N+1}\right). \tag{S61}$$

Rearrangement and factoring gives

$$\frac{r\nu}{\alpha(1-\delta)}\left[1 - (1-\delta)\frac{k}{N} - \delta\left(\frac{k}{N+1} + \frac{k}{N(N+1)}\right)\right]. \tag{S62}$$

Application of Eq. S59 and evaluation ultimately gives

$$\frac{r\nu}{\alpha(1-\delta)}\left(1 - \frac{k}{N}\right), \tag{S63}$$

which is again $V(x)$ for $x > x_I$. □

A comparison of the finite-memory value of Eq. S48 to the infinite-memory value given by Eq. S56 and its convergence is depicted by Fig. 2.

### S6 Optimal memory adaptation in changing environments

In this section, we consider a cell optimally navigating a fluctuating environment but also capable of varying its memory capacity. We consider a switch between two fluctuating environmental states: $L$ with a corresponding rate of '$H$-type' signal arrival given by $p_L$, and $H$ with $H$-type signal arrival rate $p_H$. We assume that $p_L < p_I < p_H$ so that the optimal growth strategy changes upon state switching. At issue is understanding the decision-making that may lead to a optimized growth dynamics. We will consider two rules for determining memory size, based either on the cell's empirical estimation of variance, or based on the proximity of the estimated $\pi$ to $p_I$.

#### S6.1 Principle of variance minimization

We denote by $\varepsilon_H \equiv p_H - p_I$ and $\varepsilon_L \equiv p_I - p_L$ the relative distances between state probabilities and the indifference probability. We assume that the population spends $M$ periods in each state before switching, and may possess a memory capacity of $1 \leq N \leq M$. Our goal is to identify the optimal memory $N^*$, $1 \leq N^* \leq M$. We refer to the population's *estimation* of the state as $E$, with the *true* state as $T$, with $E, T \in \{L, H\}$. In a manner similar to Eq. S33, the probability of *incorrectly estimating* the current period given the random sum of $K_\ell$ $H$-type signals out of $N$ total trials, $\ell$ of which occur after switching to state $H$, is given by the one-sided Chebyshev estimates:

$$\mathbb{P}\left(E = H \mid T = L\right) = \mathbb{P}\left(\frac{K_\ell}{N} - p_L > p_I - p_L\right) \leq \frac{\sigma^2}{\sigma^2 + \varepsilon_L^2} \tag{S64}$$

and

$$\mathbb{P}\left(E = L \mid T = H\right) = \mathbb{P}\left(p_H - \frac{K_\ell}{N} > p_H - p_I\right) \leq \frac{\sigma^2}{\sigma^2 + \varepsilon_H^2} \tag{S65}$$

with $\sigma^2 \equiv \mathbb{V}\mathrm{ar}\left(K_\ell/N\right)$. $N_\ell$ is comprised of a sum of two binomial random variables, one representing trials obtained in the $L$ state and the other trials from the $H$ state. In particular, the more sampling that occurs (for larger $\ell$), the more that the average signal reflects the new $H$ state than the old. We may thus represent $K_\ell$ by

$$K_\ell = X_{L,N-\ell} + X_{H,\ell} \tag{S66}$$

with $X_{L,N-\ell} \sim \mathrm{Binom}(N-\ell, p_L)$ representing the number of events after $\ell$ new observations that were derived from the previous ($L$) state. Similarly, $X_{H,\ell} \sim \mathrm{Binom}(\ell, p_H)$ represents the number of events sampled from the updated ($H$) state.

Then, for $x_i \equiv p_i(1-p_i)$, $i \in \{L, H\}$, the observed estimate has variance given by

$$\begin{aligned}\sigma_{\ell,N}^2 = \mathbb{V}\mathrm{ar}\left(\frac{1}{N}\left[X_{L,N-\ell} + X_{H,\ell}\right]\right) &= \frac{1}{N^2}\Big[(N-\ell)p_L(1-p_L) + \ell p_H(1-p_H)\Big] \\ &= \frac{1}{N}\Big[x_L + \frac{\ell}{N}\Big(x_H - x_L\Big)\Big].\end{aligned} \tag{S67}$$

Clearly, this variance linearly interpolates between the limiting case of $N$ $L$-sample draws, and $N$ $H$-sample draws. Now, by Eq. S65, a simple optimal memory scheme may be obtained:

$$\underset{1\leq N\leq M}{\arg\min} \sum_{\ell=1}^{M} \frac{\sigma_{\ell,N}^2}{\sigma_{\ell,N}^2 + \varepsilon_H^2} = \underset{1\leq N\leq M}{\arg\min}\left\{\sum_{\ell=1}^{N} \frac{\sigma_{\ell,N}^2}{\sigma_{\ell,N}^2 + \varepsilon_H^2} + (M-N)\left(\frac{\sigma_{N,N}^2}{\sigma_{N,N}^2 + \varepsilon_H^2}\right)\right\}. \tag{S68}$$

**Proposition 2.** *The optimal memory scheme that solves Eq. S68 is given by minimizing $N$ if $p_L(1-p_L) > p_H(1-p_H)$ and maximizing $N$ if $p_L(1-p_L) < p_H(1-p_H)$.*

*Proof.* Put $S_{\ell,N} \equiv \sigma^2_{\ell,N}/(\sigma^2_{\ell,N}+\varepsilon^2_H)$. By S67 we have, for the derivatives of $\sigma^2_{\ell,N}$:

$$\frac{\partial S^2_{\ell,N}}{\partial \ell} = \varepsilon^2_H \frac{(x_H - x_L)}{\left[x_L + \frac{l}{N}(x_H - x_L) + N\varepsilon^2_H\right]^2}; \tag{S69}$$

$$\frac{\partial S^2_{\ell,N}}{\partial N} = -\varepsilon^2_H \frac{\frac{2\ell}{N}\big(x_L - x_H\big) - x_L}{\left[x_L + \frac{l}{N}(x_H - x_L) + N\varepsilon^2_H\right]^2}. \tag{S70}$$

**Case I:** $x_H > x_L$. In this case $S^2_{\ell,N}$ is an increasing sequence in $\ell$, so $S^2_{\ell,N} \le S^2_{N,N}$ for each $N$. Since $S^2_{\ell,N}$ is decreasing in $N$, Eq. S68 is minimized at $N^* = M$.

**Case II:** $x_H < x_L$. Here, it is useful to interpret $N\sigma^2_{\ell,N}$ as a convex combination of $x_H$ and $x_L$:

$$N\sigma^2_{\ell,N} = x_L + \frac{\ell}{N}\Big(x_H - x_L\Big) = \frac{\ell}{N}x_H + \Big(1 - \frac{\ell}{N}\Big)x_L, \quad \text{for } 0 \le \ell \le N. \tag{S71}$$

$x_H < x_L$ together with the monotonicity of the function $f(x) = x/(x+y)$ implies

$$\frac{x_H}{x_H + N\varepsilon^2_H} < \frac{x_L + \frac{\ell}{N}\Big(x_H - x_L\Big)}{x_L + \frac{\ell}{N}\Big(x_H - x_L\Big) + N\varepsilon^2_H} \tag{S72}$$

for each $N$, with $1 \le \ell \le N$. The left-hand term of Eq. S72 gives

$$M\left(\frac{x_H}{x_H + N\varepsilon^2_H}\right) = N\left(\frac{x_H}{x_H + N\varepsilon^2_H}\right) + (M-N)\left(\frac{x_H}{x_H + N\varepsilon^2_H}\right). \tag{S73}$$

Thus, applying inequality of Eq. S72 for each $N$,

$$M\left(\frac{x_H}{x_H + N\varepsilon^2_H}\right) \le \sum_{\ell=1}^{N} \frac{x_L + \frac{\ell}{N}\Big(x_H - x_L\Big)}{x_L + \frac{\ell}{N}\Big(x_H - x_L\Big) + \varepsilon^2_H} \; + \; (M-N)\left(\frac{x_H}{x_H + \varepsilon^2_H}\right) \tag{S74}$$

The right hand side is none other than the sum of $\sigma^2_{\ell,N}/(\sigma^2_{\ell,N}+\varepsilon^2_H)$ terms, which implies that the minimizer of Eq. S68 is $N^* = 1$. □

The above result offers one postulate for the behavior of a cell determining the optimal memory when navigating a new state: If the variance in individual signal arrivals experienced by the population in the new environmental state is greater than in the previous state, then the advantage that a large memory enjoys outweighs the cost of momentum associated with averaging the prior state in the long run. Otherwise, if the variance of the new state is smaller than the previous state, then the strategy that opts to forget all past information in the average except for the most recent signal is preferred. We note an identical conclusion for the the case where the roles of $H$ and $L$ states are reversed. This result is independent of the current variance or relative positions of $p_H$ and $p_L$ to $p_I$.

To implement this, we can consider a cell that assesses its observed sample variance $\pi_n(1-\pi_n)$ at each period $n$, with $\pi_n = k_n/N_n$. If the sampled variance increases from one period to the next, then the memory size in the next period, $N_{n+1}$, increases to $N_n + 1$. Similarly, variance reductions result in updated a reduction in memory size by one or no change. If the variance is equal, there is no change in the memory.

Curiously, it cannot be that memory decreases if there is a tie. For if so, then under adaptive environments, lower memories occur when the sampled variance is equal, which can inappropriately drive depleted memories when they are needed (Fig. S2B). In comparing fixed vs. adaptive memory capacity systems, large initial memory sizes in constant-to-fluctuating, persistence of memory is predicted to lead decreased growth performance early on. This is followed by improvements in long-run growth as the initial memory begins to observe the fluctuating environment. We expect this case to be less likely as in the variance-based memory selection scheme, constant environments have low variance.

#### S6.2 Principle of proximity to indifference point

An alternative objective by which the cells may attempt to navigate their environment can be simply stated by the following rule: Cells leverage additional memory whenever their point-estimate of $p$ is closer to $p_I$ in order to more accurately determine the current state. In this scheme, memory evolves based on $f(d_n)$, with $d_n = |\pi_n - p_I|$. $f$ may in general be an arbitrary, non-decreasing function. For foundational understanding in the case of finite memory, we assume a linear dependence, given by rounding the following functional form:

$$f_n = N_{\max} - (N_{\max} - N_{\min})d_n, \tag{S75}$$

having incremental transitions ($\pm 1$ memory unit each period) and interpolating between a maximal and minimal memory capacity:

$$N_{n+1} = \begin{cases} \min\{N_{\max}, N_n + 1\}, & f_{n+1} > f_n; \\ N_n, & f_{n+1} = f_n; \\ \max\{N_{\min}, N_n - 1\}, & f_{n+1} < f_n. \end{cases} \tag{S76}$$

or more concisely as

$$N_{n+1} = N_n + I_{[d_{n+1}<d_n]\&[N_n<N_{\max}]} - I_{[d_{n+1}>d_n]\&[N_n>N_{\min}]}. \tag{S77}$$

where

$$I_E = \begin{cases} 1, & E; \\ 0, & E^c. \end{cases} \tag{S78}$$

We can illustrate the temporal evolution of adaptive memory relative to what would be expected from a known environmental landscape, $N_\infty$, which is given by setting $\pi = p$ above in Eq. S75. Plots of adaptive memory behavior governed by proximity are given in Figs. 4B-C, S3. From these, it is clear that memory adaptation enables cells to ultimately achieve the highest growth potential observed across their fixed-memory counter parts.

### S7 Application to cellular memory in a fluctuating environment

Our goal here is to apply the finite fixed and adaptive memory models to compare predicted growth strategies for cell populations encountering a constant versus a rapidly-fluctuating environment to those observed experimentally. A surprising recent result was the observation of a distinct growth phenotype observed for cells undergoing rapid fluctuations that is lower than the corresponding averaged nutrient level of their constant-environment counterparts. Toward that end, our framework below will be applied to support the probabilistic basis for this finding.

#### S7.1 Physiologic adaptation to variable nutrient environment

We consider a cell faced with either a low ($L$) depleted nutrient environment, or a high ($H$) enriched nutrient environment. Cells may correspondingly adapt either a phenotype that is flexible and advantageous in the $H$-environment, denoted by $S_{High}$, or a phenotype that is preferred in the $L$-environment, denoted by $S_{Low}$. The corresponding growth rate per unit time under the $S_{Low}$ and $S_{High}$ phenotypes can be written respectively as

$$R_{Low,n} = \beta r_L(1 - H_n), \qquad R_{High,n} = r_L(1 - H_n) + r_H H_n, \tag{S79}$$

with $\mathbb{P}(H_n = 1) = p$ characterizing the likelihood of nutrient rich conditions and $\beta > 1$ representing the relative gain in efficiency for committing to growing under low nutrient availability. This efficiency $\beta$ is directly comparable to the original setup that considered costly environments with the flexible $S_A$ phenotype; here, we consider $\beta = 1/\alpha$ for $\alpha < 1$. As before, the $S_{Low}$-phenotype is preferred whenever $p < p_I$, and the $S_{High}$-phenotype is preferred whenever $p > p_I$. In this case,

$$p_I = \frac{(\beta - 1)r_L}{(\beta - 1)r_L + r_H}. \tag{S80}$$

Motivated by solving the above for $\beta$ evaluated the point of indifference,

$$\beta = 1 + \frac{r_H}{r_L}\left(\frac{p_I}{1 - p_I}\right). \tag{S81}$$

we will subsequently discuss $\beta$ relative to a characteristic value $\beta_c = 1 + r_H/r_L$, at which $p_I = 1/2$.

#### S7.2 Fixed memory momentum and resolvability

**Momentum:** Systems with large memories that persist in one environment before switching to another have an ingrained momentum associated with the old environment that takes longer before their memory may begin to detect the new environment. We characterize this mathematically in the case where a population navigates a constant environment (with $p_0 = 0$) to a fluctuating one (with $p_1 = p$). We compare the system's estimated environment $K_\ell/N$ with memory capacity $N$ after spending $\ell$ moments in the new fluctuating environment with the critical indifference environment $p_I$. The experienced environment is a sum of constant and variable terms, each Binomially-distributed with either $N - \ell$ or $\ell$ occurrences and 0 or $p$ success probability, respectively:

$$K_\ell = X_{C,N-\ell} + X_{V,\ell}. \tag{S82}$$

Thus, in this case, $X_{C,N-\ell} = 0$, and $X_{V,\ell} \sim \text{Binom}(\ell, p)$. The point at which the system's phenotypic choice begins to reflect the new environment is given in expectation whenever $\mathbb{E}[K_\ell]/N = p_I$, which occurs at

$$\ell_{Crit} = \frac{p_I}{p}N. \tag{S83}$$

$\ell_{Crit}$ therefore quantifies the time prior to 'unlearning' a constant state in a new variable environment and scales directly with memory size.

**Resolvability:** While larger memory sizes take longer to reliably estimate new environments, their estimate of those environments are more accurate. In a manner similar to that of Eqs. S64,S65, this particular case may be further simplified. For a system initially navigating a constant 0 state to a fluctuating state such that $p > p_I$, the probability of mis-estimating the current state is given by

$$\mathbb{P}\left(K_\ell/N \leq p_I\right) = \mathbb{P}\left(X_{V,\ell} \leq p_I N\right) = F_\ell(p_I N), \tag{S84}$$

where $F_\ell(x)$ is the cumulative distribution function for a Binomial$(\ell, p)$ random variable. The probability of correctly estimating the state is therefore $1 - F_k(p_I N)$. Assuming that the growth rate for the constant (resp. fluctuating) environment is given by $R_L$ (resp. $R_V$), and $R_L < R_V$, the growth rate may therefore be given exactly by

$$R = R_F\left[1 - F_k(p_I N)\right] + R_L F_k(p_I N). \tag{S85}$$

Plots of the optimal growth potential for a variety of memory sizes and efficiencies are given in Figs. 4, S4, from which we find excellent agreement with simulations in the maximal ultimate growth rates given in Eq. S85 and the critical transition times given in Eq. S83. When considering a moving window of averaged growth rates over successive times, these results are in agreement with empirical observations that demonstrate improvements in the growth rate with successive fluctuations [1] in nutrient-high environments, with an inflection occurring after a small (roughly 4) number of switches in the fluctuating environment. Moreover, the empirical growth curves appear to perform better initially in fluctuating-high environments, and worse initially in fluctuating-low environments, suggesting that in this experimental setup, $p = 1/2$ is larger than $p_I$. We further consider a dynamic variance- and proximity-driven memory update schemes as described above (Figs. S2,S3).

#### S7.3 Jensen's Inequality Discussion

Another important empirical observation concerns the fact that cells in rapidly fluctuating nutritional environments grow at slower rates than cells growing in a corresponding constant environment that averages out the total nutrient content [1]. Significantly, this observation is demonstrated to persist even after accounting for the convexity of the growth-versus-nutrient curve. One principal advantage of our modeling approach is that we have assumed throughout that the growth potential being studied represents the amount of nutrient resources $R$ available for growth, with an assumed one-to-one correspondence to growth rate $g(R)$. Approaching the problem this way enables us to avoid the complication of quantifying the expected growth in fluctuating environments, $\mathbb{E}[g(R)]$, compared to the growth rate of the corresponding averaged constant environment, $g(\mathbb{E}[R])$, for which Jensen's inequality would give $\mathbb{E}[g(R)] \leq g(\mathbb{E}[R])$. In general, obtaining an explicit formula for the deficit described by Jensen's inequality can be difficult, and so this makes quantifying any further reductions in growth challenging. Jensen's inequality holds at equality in our case since we are directly considering linear combinations of random variables, thus enabling a direct quantification of these deficits.

We observed previously that inference can result in improper estimation of the observed environment and subsequent phenotypic mismatch, and that this mismatch can be mitigated by systems with larger memories. We henceforth discuss another intrinsic misestimation risk relative to constant environments that cannot be mitigated by larger memories. One key conclusion from the above empirical estimates is that cells pursue distinct phenotypic growth strategies when faced with constant versus fluctuating environments. We apply our framework to compare the latter case with cells growing in corresponding constant environments. In the constant-environment case, cells maximally adapt to acquiring nutrients in a manner proportional to those available. The average growth potential can therefore be represented as a convex combination $p$ of deterministic environments representing the extreme $H$- and $L$-type environments (assuming environmental landscapes of $H_n = 1$ and $H_n = 0$, respectively). Thus, growth in the constant average environment is given by:

$$R_{avg} = (1-p)\beta r_L + p r_H. \tag{S86}$$

This growth rate corresponds to the constant-environment analog of the random growth policy under fluctuating environments having parameter $p$:

$$R_{\text{rand}} = \begin{cases} (1-p)\beta r_L, & S_{Low}\text{-phenotype;} \\ (1-p) r_L + p r_H, & S_{High}\text{-phenotype.} \end{cases} \tag{S87}$$

Now, in the best case, the cell correctly matches its phenotype to the environment $S_{Low}$ when $p < p_I$, $S_{High}$ when $p > p_I$. Calculating the difference between average and random values for this case gives the *intrinsic growth deficit*, $D_I$:

$$D_I = \begin{cases} p r_H, & p < p_I; \\ \dfrac{(\beta-1) r_L r_H}{(\beta-1) r_L + r_H}, & p = p_I; \\ (1-p)(\beta-1) r_L, & p > p_I. \end{cases} \tag{S88}$$

Eq. S88 is piecewise-linear, and maximized at $p = p_I$. $D_I$ represents the ($N$-independent) minimal possible deficit assuming a proper phenotypic choice is made and is a universal property of systems that select phenotypic choices based on past environmental inference. This results from the fact that stochastic environments will inevitably have fluctuations that shock the cell population, despite its selection of the correct phenotype maximizing expected long-term growth. In the event that the phenotype is mis-matched to the environment, there is an additional *extrinsic growth deficit*, $D_E$, which can be represented by

$$D_E = |(1-p)(\beta-1) r_L - p r_H| \, I_{miss}. \tag{S89}$$

Here, $\mathbb{P}(I_{miss}=1) = p_{miss}$ describes the probability of phenotypic mismatch. $p_{miss}$ is related to the cumulative distribution function $F_K(k)$ for $K \sim \text{Binomial}(N,p)$ (see Sec. S7.2). This occurs with probability $1 - F_K(p_I N)$ whenever $p < p_I$, $F_K(p_I N)$ when $p > p_I$. A normal approximation gives

$$p_{\text{miss}} \approx \begin{cases} 1 - \Phi\left((p_I - p)\sqrt{\frac{N}{p(1-p)}}\right), & p < p_I; \\ \frac{1}{2}, & p = p_I; \\ \Phi\left((p_I - p)\sqrt{\frac{N}{p(1-p)}}\right), & p > p_I. \end{cases} \tag{S90}$$

Together, we have the total deficit given by the sum of the deterministic $D_I$ and random $D_E$ components

$$D_{\text{tot}} = D_I + D_E. \tag{S91}$$

and we can from this calculate the mean deficit as functions of $\beta$ and $p$. Our results demonstrate that a growth deficit will always exist across all environmental and cell-specific parameters. The intrinsic deficit is maximized at the point of environmental indifference $p_I$, which converges to the total deficit whenever the memory term becomes large. For smaller memory terms, there is enhancement of the deficit owing to phenotypic mismatch. These results are quantified in Figs. 5B-C, S6. Intrinsic deficits for larger memory sizes are depicted in Fig. 5D, while the influence of memory and an example of the discrepancy between growth deficits deviating away from $p_I$ are given in Fig. 5E-F.

#### S8 Supplementary Figures

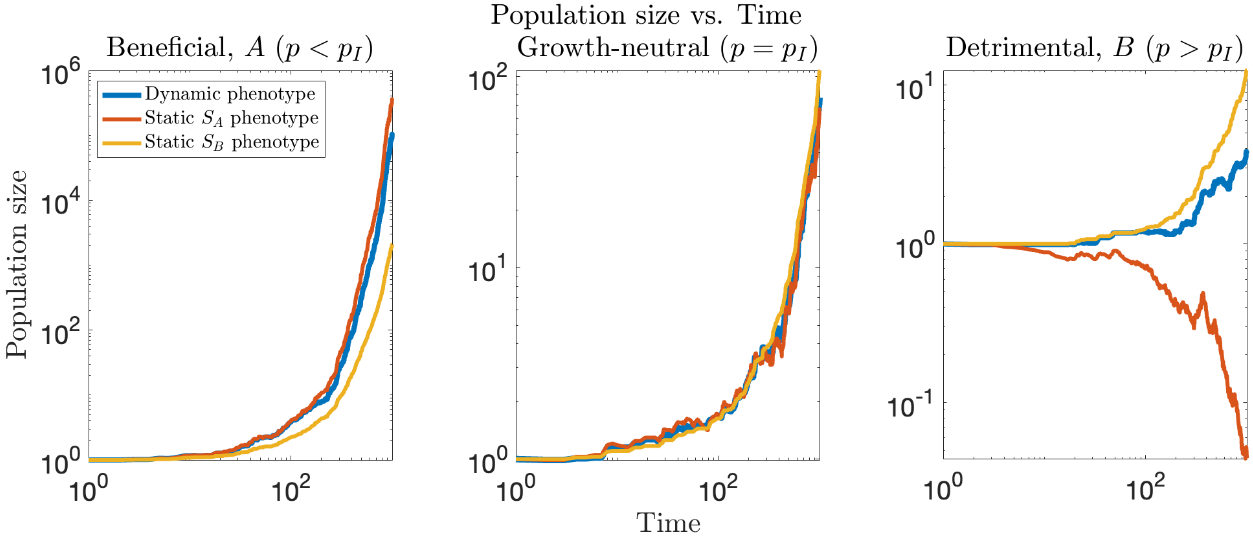


Figure S1: Growth in fluctuating environments. Representative stochastic trajectories of the population size are depicted for cells navigating type-$A$ (left), netural (middle), and type-$B$ (right) environments. Adaptive systems (blue) capable of phenotypic switching have lower growth potential than static phenotypes (red, yellow) matched to the proper environment, but outperform those phenotypes whenever the environment switched. All phenotypic strategies collapse in a neutral environment (In all cases, $r\nu$=2, $c\lambda$=1, $\alpha$=2, $\delta$=0.9, $N$=20).

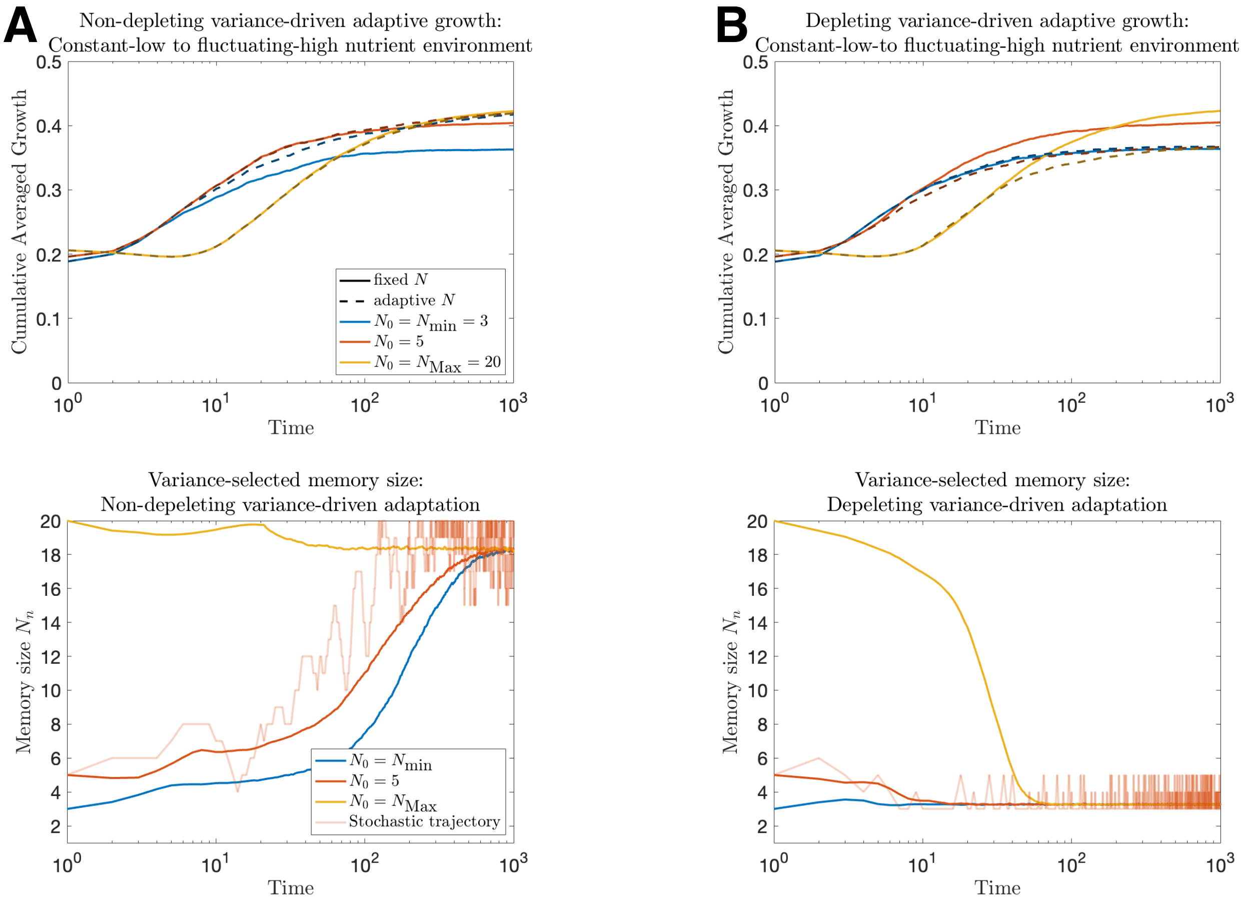


Figure S2: Dynamics of fixed and adaptive memory systems navigating constant-low to fluctuating-high nutrient environments. (A) Adaptive cells navigate their environment by changing their memory capacity based on the relative variance between successive environments (Sec. S6.1) assuming that ties do not result in reduced memory capacities. In this case, memory capacity stochastically increases to improve decision-making that leads to improved growth in adaptive systems relative to their fixed, low-memory counterparts. (B) It is important that the memory update scheme distinguish reductions in variance from stationary values, as this would drive adaptive cells toward lower memories resulting in a loss of growth (in all cases, $r_H = 1$, $r_L = 0.05$, $N_{min} = 3$, $N_{Max} = 20$, $\beta = \beta_{Crit}/5$ giving $p_I = 0.14$, $p = 0.4$. Constant-low nutrient environments were given by the deterministic sequence $L_n = 0$, while fluctuating-high nutrient environments by $\mathbb{P}(L_n = H) = p$, with $p > p_I$. For each initial memory condition $N_0$ and update scheme considered, reported values are averaged over $10^3$ iterates).

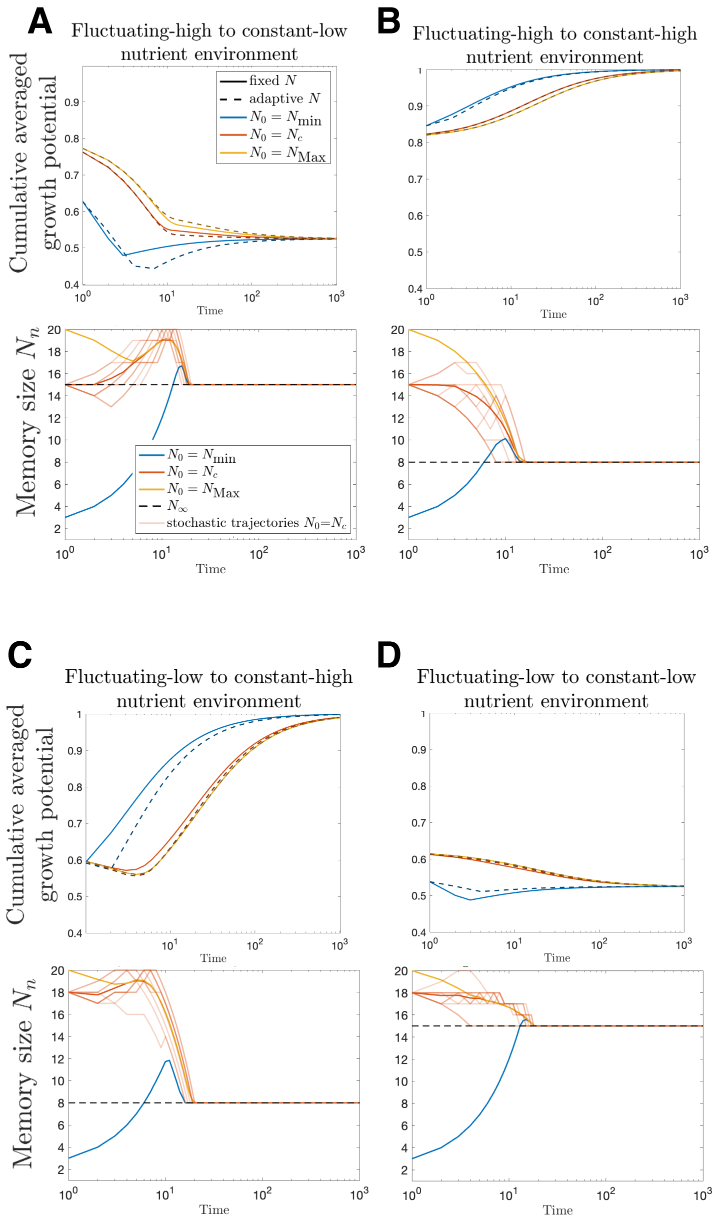


Figure S3: Dynamics of fixed and adaptive memory systems navigating fluctuating to constant environments. Adaptive cells navigate their environment by changing their memory capacity based on the proximity of the estimated environment to the critical indifference environment $p_I$ (Sec. S6.2) are compared to fixed-memory counterparts when navigating (A) fluctuating-high to constant-low, (B) fluctuating-high to constant-high, (C) fluctuating-low to constant-high, and (D) fluctuating-low to constant-low nutrient environments. Constant-low (resp. high) nutrient environments were given by the deterministic sequence $L_n = 0$ (resp. $L_n = 1$), while fluctuating-high (resp. low) nutrient environments by $\mathbb{P}(L_n = H) = p$, with $p > p_I$ (resp. $p < p_I$). The theoretical prediction of $N_c$ is given by evaluating Eq. S75 with the corresponding $p$ and $p_I$. For A,B $p = 0.6$ describes the fluctuating-high environment, while for C,D $p = 0.20$ describes the fluctuating-low environment. (in all cases, $r_H = 1$, $r_L = 0.05$, $N_{min} = 3$, $N_{Max} = 20$, $\beta = \beta_C/2$, giving $p_I = 0.32$. For each initial memory condition $N_0$ and update scheme considered, reported values are averaged over $10^3$ iterates).

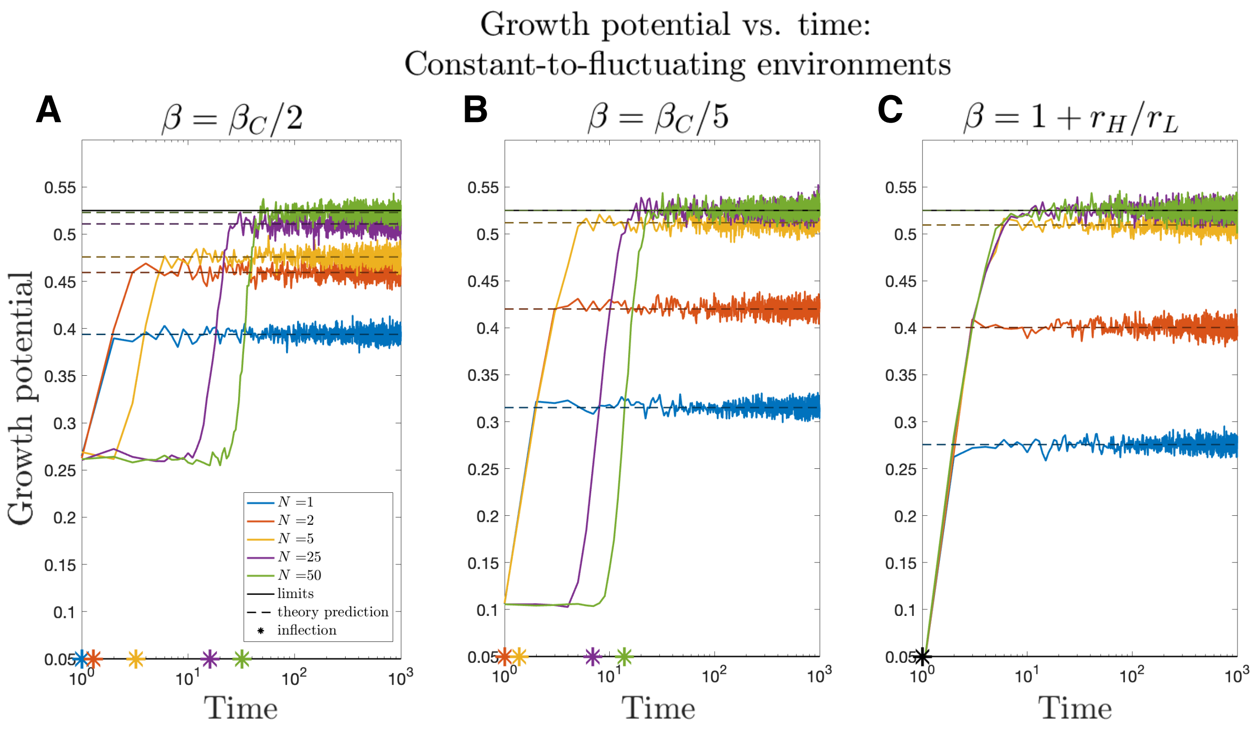


Figure S4: The effects of memory size on maximal realizable growth and adjustment time for systems navigating constant-low to fluctuating-high nutrient environments. Growth potential as a function of time is depicted for a variety of fixed-memory systems navigating the onset of rapid nutrient fluctuations, assuming (A) significant ($\beta = \beta_C/2$), (B) intermediate ($\beta = \beta_C/5$), or (C) mild ($\beta = 1 + r_L$) growth efficiencies of the $S_{Low}$ phenotype. The theoretically predicted long-run growth potential and inflection points are given by dashed horizontal lines and stars, respectively (in each case, $r_L = 0.05$, $r_H = 1$ giving $\beta_C = 1 = r_H/r_L$, $p = 0.6$).

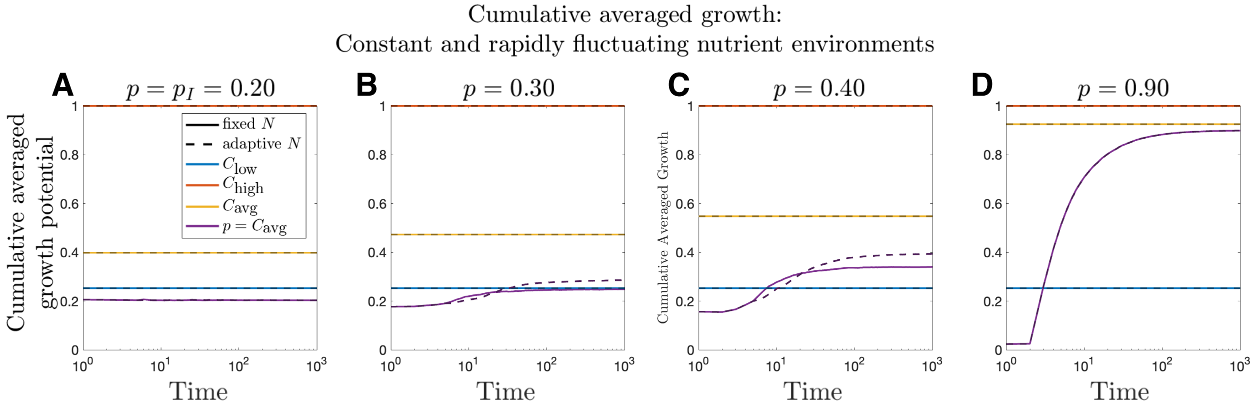


Figure S5: Growth dynamics for cells in fluctuating nutrient environments and comparable constant environments. Cumulative averaged growth potentials are given as a function of time for cells with fixed (solid) and adaptive (dashed) memory in either nutrient rich (red), depleted (blue), averaged (yellow), or fluctuating (purple) environments. Convex combinations of mean nutrient rich and poor environments were taken to match the environmental parameter $p$ governing the probability of a rich nutrient environment. For a variety of increasing environmental values of $p$, growth in constant average environments surpass the corresponding fluctuating environments of having equal likelihoods of nutrient rich signal (In all cases, $r_L = 0.01$, $r_H = 1$, $\beta_C = 1 + r_H/r_L$, $\beta = \beta_C/4$, giving $p_I = 0.20$; $10^3$ stochastic simulations were evaluated for each simulation over time).

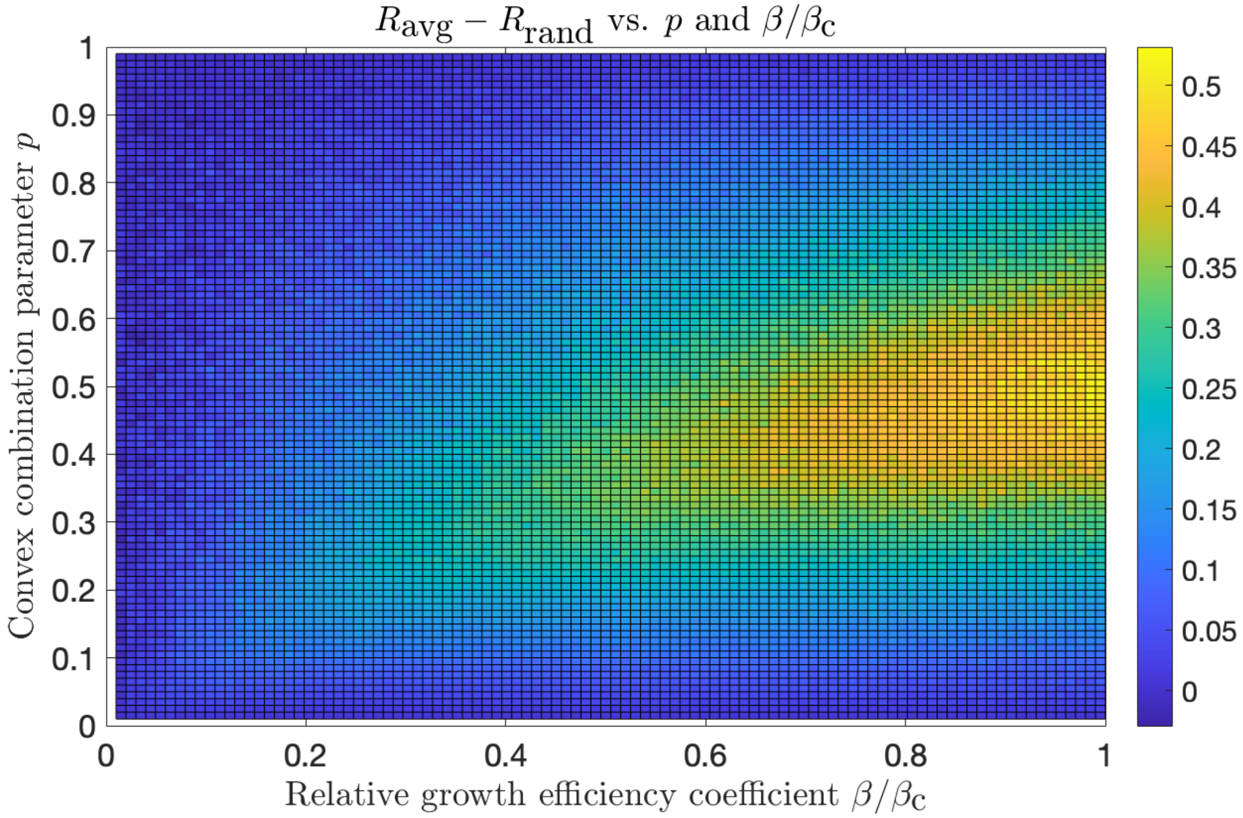


Figure S6: Growth deficits always occur in fluctuating environments. The difference in the simulated long-term growth potential between constant environments and their corresponding fluctuating environments is given across all environmental parameters $p$ and allowable growth coefficients $\beta$ (point-wise difference of Fig. 5C and 5B; $r_L = 0.01$, $r_H = 1$, $\beta_C = 1 + r_H/r_L$).
